# Structural Differences In 3C-like protease (Mpro) From SARS-CoV and SARS-CoV-2: Molecular Insights For Drug Repurposing Against COVID-19 Revealed by Molecular Dynamics Simulations

**DOI:** 10.1101/2021.08.11.455903

**Authors:** Meet Parmar, Ritik Thumar, Bhumi Patel, Mohd Athar, Prakash C. Jha, Dhaval Patel

**Affiliations:** Department of Biological Sciences and Biotechnology, Indian Institute of Advanced Research, Gujarat-382426, India; Center for Chemical Biology and Therapeutics, InStem, Bangalore-560065, Karnataka, India; School of Applied Material Sciences, Central University of Gujarat, Gandhinagar-382030, Gujarat, India

## Abstract

A recent fatal outbreak of novel coronavirus SARS-CoV-2, identified preliminary as a causative agent for series of unusual pneumonia cases in Wuhan city, China has infected more than 20 million individuals with more than 4 million mortalities. Since, the infection crossed geographical barriers, the WHO permanently named the causing disease as COVID-2019 by declaring it a pandemic situation. SARS-CoV-2 is an enveloped single-stranded RNA virus causing a wide range of pathological conditions from common cold symptoms to pneumonia and fatal severe respiratory syndrome. Genome sequencing of SARS-CoV-2 has revealed 96% identity to the bat coronavirus and 79.6% sequence identity to the previous SARS-CoV. The main protease (known as 3C-like proteinase/ Mpro) plays a vital role during the infection with the processing of replicase polyprotein thus offering an attractive target for therapeutic interventions. SARS-CoV and SARS-CoV-2 Mpro shares 97% sequence identity, with 12 variable residues but none of them present in the catalytic and substrate binding site. With the high level of sequence and structural similarity and absence of any drug/vaccine against SARS-CoV-2, drug repurposing against Mpro is an effective strategy to combat COVID-19. Here, we report a detailed comparison of SARS-CoV-2 Mpro with SARS-CoV Mpro using molecular dynamics simulations to assess the impact of 12 divergent residues on the molecular microenvironment of Mpro. A structural comparison and analysis is made on how these variable residues affects the intra-molecular interactions between key residues in the monomer and biologically active dimer form of Mpro. The present MD simulations study concluded the change in microenvironment of active-site residues at the entrance (T25, T26, M49 and Q189), near the catalytic region (F140, H163, H164, M165 and H172) and other residues in substrate binding site (V35T, N65S, K88R and N180K) due to 12 mutation incorporated in the SARS-CoV-2 Mpro. It is also evident that SARS-CoV-2 dimer is more stable and less flexible state compared to monomer which may be due to these variable residues, mainly F140, E166 and H172 which are involved in dimerization. This also warrants a need for inhibitor design considering the more stable dimer form. The mutation accumulated in SARS-CoV-2 Mpro indirectly reconfigures the key molecular networks around the active site conferring a potential change in SARS-CoV-2, thus posing a challenge in drug repurposing SARS drugs for COVID-19. The new networks and changes in microenvironment identified by our work might guide attempts needed for repurposing and identification of new Mpro inhibitors.

## Introduction

The year 2019 ended with a fatal outbreak of a novel coronavirus (SARS-CoV-2) identified preliminary as a causative agent for series of unusual pneumonia cases in Wuhan city, Hubei province of China [1,2]. As the cases were reported sporadically, the WHO announced a Public Health Emergency of International Concern (PHEIC) for the 2019-nCoV outbreak in January 2020 (WHO.int). Since the infection crossed geographical barriers, the WHO permanently named the 2019-nCoV pathogen as SARS-CoV-2 and the causing disease as coronavirus disease 2019 (COVID-2019) by declaring it a pandemic situation in March 2020 (WHO.int). As per the latest situation report (8^th^ August 2021) of WHO, more than 20 million confirmed cases with more than 4 million death are reported globally affecting 208 countries in the world (WHO.int). The disease caused by SARS-CoV-2 presents vast pathophysiological symptoms including fever, coughing and shortness of breath in common cases whereas pneumonia, kidney failure and severe acute respiratory failure in severe cases [2,3].

The causative agent for SARS-CoV-2, coronavirus belongs to subgenus Sarbecovirus, Coronaviridae family and Orthocoronavirinae subfamily. It is a β-coronavirus of 2B (β-coronavirus) group till date not reported zoonotic unlike earlier coronaviruses [4,5]. SARS-CoV-2 is an enveloped single-stranded RNA virus (+ve ssRNA) with a genome size of with 29.9 kb genome that spreads widely among humans and other mammals, causing a wide range of infections from common cold symptoms to fatal diseases, such as severe respiratory syndrome [6,7]. Genome sequencing of SARS-CoV-2 has revealed 96% identity to the bat coronavirus and 79.6% sequence identity to SARS-CoV [3]. Other known members of β-coronavirus which are pathogenic to humans are SARS-CoV ("severe acute respiratory syndrome"), the causative agent for the outbreak in 2002-2003 [8] and MERS-CoV ("Middle East respiratory syndrome") outbreak in 2012 [9] with a high mortality rate of 9.6% for SARS-CoV and 34.4% for MERS-CoV. Despite being genetically distinct from SARS-CoV, the proteome is quite similar but have distinct features of which two major proteins have been identified for pathogenicity: the non-structural proteins (Nsps) and structural proteins. The SARS-CoV-2 virus genome has 15 ORFs: ORF 1a (encodes non-structural proteins - nsp1 to nsp11), ORF 1b (encodes non-structural proteins (nsp12 to nsp16), ORF S (encodes a Spike protein), ORF 3a, 3b, 6, 7a, 7b, 8, 9a, 9b and 10 (encodes accessory protein), ORF E (encodes envelope protein, virus assembly and morphogenesis proteins), ORF M (encodes membrane protein and virus assembly proteins) and ORF N (encodes nucleocapsid protein) (Lu et al., 2020; Wu et al., 2020). With the aid of ribosomal frame-shifting during translation, the replicase gene of SARS-CoV-2 encodes two over-lapping translation products, polyproteins 1a and 1ab (pp1a and pp1ab) [12]. Each of these polyproteins is then cleaved by main protease (Mpro) and PLpro (Papain-like Protease) of SARS-CoV-2 to release 11 (pp 1a) and 5 (pp 1ab) functional proteins necessary for viral replication.

As viral proteases play a vital role in processing the polyproteins that are translated from the viral RNA, inhibitors designed against these proteases may prove effective in blocking the replications of coronaviruses thus making them attractive target for drug discovery [6]. Viral proteases have been suggested to have relationships with the mechanism of infection and pathogenicity of SARS-CoV-2 [13]. The non-structural protein 5 (Nsp5) is the main protease (Mpro) of SARS-CoV-2, a chymotrypsin-like cysteine protease also known as "3C-like protease" (3CLpro) necessary for viral replication, structural assembly and pathogenicity [14]. The Mpro (3CLpro) in particular cleaves at least at 11 sites during proteolytic processing of polyproteins pp1a and pp1ab. The recognition sequence at most sites is Leu-Gln↓(Ser, Ala, Gly) (↓ marks the cleavage site) [15,16]. Mpro protease of SARS-CoV-2 is classified as a chymotrypsin-like cysteine protease (3CLpro) with a molecular mass of ~33.8KDa [17]. The recent availability of SARS-CoV2 Mpro structure reveals the presence of three domains; the domains I (residues 1–101) and II (residues 102–184) made up by antiparallel β-barrel structures (13 β-strands) in a chymotrypsin-like fold responsible for catalysis and the α-helical domain III (residues 201–306) (5 α-helices) is required for the enzymatic activity [16,17]. A long loop (residues 185-200) connects domains I, II with III. The substrate-binding site is located in a cleft between these domains I and II comprising of a Cys-His dyad with an overall structure very similar to SARS-CoV Mpro as well as other SARS family Mpro (MERS-Mpro, HKU5-Mpro, HKU4-Mpro and SARS-Mpro [17]. The structures SARS-CoV-2 Mpro and SARS-CoV Mpro are quite similar, the main difference being the surface of the proteins, where 12 different amino acids are located. Since the first structure of SARS-CoV was available and more recently the structure of SARS-CoV-2 was solved, numerous efforts have been made to examine it as a potential target to prevent the spread of infection by inhibiting viral polyprotein cleavage through blocking active sites of the protein [15,16]. Our present work attempting to identify the sequence and structural differences between SARS-CoV Mpro and SARS-CoV-2 Mpro as well as the differences in dynamics of both the Mpro was investigated using Molecular Dynamics approach. The present findings may help researchers to identify new compounds/inhibitors against SARS-CoV-2 Mpro as well as collectively knowledge on the structural dynamics vis-a-vis between two Mpro for the repurposing of existing anti-viral protease inhibitors.

## Materials and Methods

### Sequence analysis

The 3C-like Mpro sequence in FASTA format was downloaded from NCBI for the SARS-CoV-2 (PDB ID: 6M03), SARS-CoV (PDB ID: 2DUC), MERS-CoV (PDB ID: 5C3N), Bat-CoV-RaTG13 (NCBI: QHR63299.1), four other human coronaviruses; HCoV-HKU (PDB ID: 3D23), HCoV-OC43 (UniProt: P0C6X6), HCoV-NL63 (PDB ID: 3TLO), and HCoV-229E (PDB ID: 2ZU2), which have been known to cause SARS in human. The SARS-CoV-2 was also checked individually with other Mpro sequences using BLAST to identify pair-wise sequence identity. Multiple sequence alignment was performed using ClustalW to identify conserved residues followed by ESPript3 for structure-based sequence alignment using SARS-CoV-2 structure as a reference. Physicochemical parameters of all CoV 3CLpro including isoelectric point, instability index, grand average of hydropathicity (GRAVY), and amino acid composition were computed using the ProtParam tool of ExPASy.

### Structure comparison of Mpro from SARS-CoV and SARS-CoV-2

Numerous studies have been going on for Mpro from SARS-CoV-2 which has led to rapid identification and deposition of 3D X-ray structures coordinates in PDB recently. The available structures of SARS-CoV-2 Mpro were downloaded from PDB for structure comparison. The SARS-CoV-2 Mpro structures have been solved in both apo- form and with ligand-bound form. For our comparative study, we used PDB ID: 6M03; (The crystal structure of COVID-19 main protease in an apo form) for SARS-CoV-2 and PDB ID: 2DUC; (Crystal structure of SARS coronavirus main proteinase (3CLPRO)) for SARS-CoV. Mpro structures from other CoVs: SARS-CoV (PDB ID: 5B6O), MERS-CoV (PDB ID: 5WKJ), BtCoV-HKU4 (PDB ID: 2YNA), HCoV-HKU1 (PDB ID: 3D23), MHV-A59-CoV (PDB ID: 6JIJ), PEDV-CoV (PDB ID: 5ZQG), FIPV-CoV (PDB ID: 5EU8), TGEV-CoV (PDB ID: 4F49), HCoV-NL63 (PDB ID: 5GWY), HCoV-229E (PDB ID: 2ZU2) and IBV-CoV (PDB ID: 2Q6D) were downloaded from PDB for structural superposition. Multiple sequence alignment was performed using ClustalW to identify conserved residues, followed by phylogenetic tree construction using MEGA6. For tree construction, Maximum Parsimony method was used followed with 1000 bootstrap iteration for statistical significance. The alignment was generated using ESPript3 using SARS-CoV-2 structure as a reference. The alignment was generated using ESPript3 using SARS-CoV-2 structure as a reference. PyMOL, Chimera and other tools were used for in-depth analysis of structural features in SARS-CoV-2 Mpro and differences with other CoV Mpro.

### Molecular Dynamics Simulations of Mpro from SARS-CoV-2 and SARS-CoV

For comparison of dynamics of Mpro from SARS-CoV and SARS-CoV-2, PDB ID: 2DUC and PDB ID: 6M03 were used as the starting conformation respectively in Molecular Dynamics Simulations (MDS). To assess the comparison between monomer and dimer, four MDS were setup: SARS-CoV_monomer, SARS-CoV_dimer, SARS-CoV-2_ monomer and SARS-CoV-2_ dimer. Each MDS setup was performed for 50ns production run as reported in our earlier studies [18,19] using GROMACS ver.2016.4 [20] with Amber99SB force-field [21]. All the MDS systems were solvated with simple point charge (SPC) water model in dodecahedron box configuration with a distance of 1 nm from all the directions of the protein and periodic boundaries. The simulations setup was neutralized by adding an equal number of counter ions (Na^+^/Cl^-^) and subjected to energy minimization using the steepest descent algorithm to remove any steric clashes and bad contacts before the actual MD run. Following minimization, equilibration with position restraint was carried out under NVT (constant number [N], constant volume [V] and constant temperature [T]) and NPT (constant number [N], constant pressure [P] and constant temperature [T]) ensemble for 1 ns each. For NVT equilibration, modified Berendsen thermostat algorithm [22] was used to maintain the system at constant volume for 100 ps and at a constant temperature of 300 K. NPT equilibration was performed at a constant pressure of 1 bar for 100 ps maintained by Parrinello-Rahman barostat [23]. For calculations of long-range electrostatic interactions, Particle Mesh Ewald approximation was applied with 1 nm cut-off [24] and computing coulomb & the van der Waals interactions and the bond length was constraint using the LINCS algorithm [25]. Final production MD was simulated for 50 ns run with default parameters. The trajectories were visualized using VMD [26] and Chimera [27]. For calculation of root mean square deviation (RMSD), root mean square fluctuation (RMSF), and hydrogen bonds (H-bonds), etc. in-built gmx commands were used in GROMACS and the plotting tool GRACE was used for the generation and visualization of the plots (http://plasma-gate.weizmann.ac.il/Grace) as reported in our previous study [28].

### Clustering of conformations for ensemble generation and Essential Dynamics

To assess the dominant conformation acquired by CoV Mpro through the entire simulations were studied by clustering analysis. The entire MDS trajectories were subjected to RMSD based clustering via ‘gmx cluster’ that explores the conformational landscape among the ensemble of protein structures. The GROMOS algorithm as described by Daura and co-workers [29], was used to determine the dominant conformation with 0.15 mm Cα RMSD cut-off. To study the global motion of SARS-CoV and SARS-CoV-2 Mpro in dimer and monomer form, Principal Component Analysis (PCA) or Essential Dynamics (ED) analysis was carried out. Similar to our earlier studies, the collective motion and essential dynamics of Cα backbone atoms were also examined for the entire simulations trajectories, as computed by using gmx covar and gmx anaeig tools [30,31]. PCA is a method that reduces the complexity of the data and results in the concerted motion in the MD simulations. These motions are essentially correlated and significant for biological functions. The set of eigenvectors and eigenvalues was computed by diagonalizing the covariance matrix. The eigenvalues represent the amplitude of the eigenvector along with the multidimensional space, while the Cα displacement along each eigenvector shows the concerted motions of the protein along each direction. FES (Free Energy Surface) (kcal/mol) was also computed for all the four MD systems considering the conformational variability in terms of ROG and RMSD took together and represented by Gibbs free energy. It represents a mapping of all possible conformations the protein adopted during the entire simulation trajectory. For computing FES, RMSD and ROG calculated to the average structure were used to ensure adequate sampling for FES calculations.

## Results and Discussion

### Sequence analysis

Multiple sequence alignment of SARS-CoV-2 3C-like Mpro was performed with its closet known homologs SARS-CoV, MERS-CoV, Bat-CoV-RaTG13, HCoV-HKU, HCoV-OC43, HCoV-NL63, and HCoV-229E to identify conserved segments in 3C-like Mpro (Figure 1). The pairwise alignment revealed that the SARS-CoV-2 3C-like Mpro is highly similar to Bat-CoV-RaTG13 (99.35%) and SARS-CoV (96.08%). It shares 50.65%, 49.02%, 48.37%, 44.30%, 41.04% sequence identity with other human coronaviruses HCoV-HKU, HCoV-OC43, HCoV-NL63, and HCoV-229E respectively (Table 1). The catalytic dyad of CYS-HIS is also conserved throughout all sequences in the Mpro family as they are very essential for enzyme activity. The physicochemical properties of SARS-CoV-2 Mpro were also in a similar range with other Mpro’s as shown in table 1. The length of Mpro from SARS-CoV-2, SARS-CoV, MERS-CoV, Bat-CoV-RaTG13 is 306 amino acids and another human coronavirus is 303 amino acids. Sequence comparison of 3C-like Mpro of SARS-CoV-2 and SARS-CoV revealed that SARS-CoV-2 has 12 divergent amino acids out of which 5 positions have residues with structurally/chemically similar amino acid substitution.

**Figure 1.**
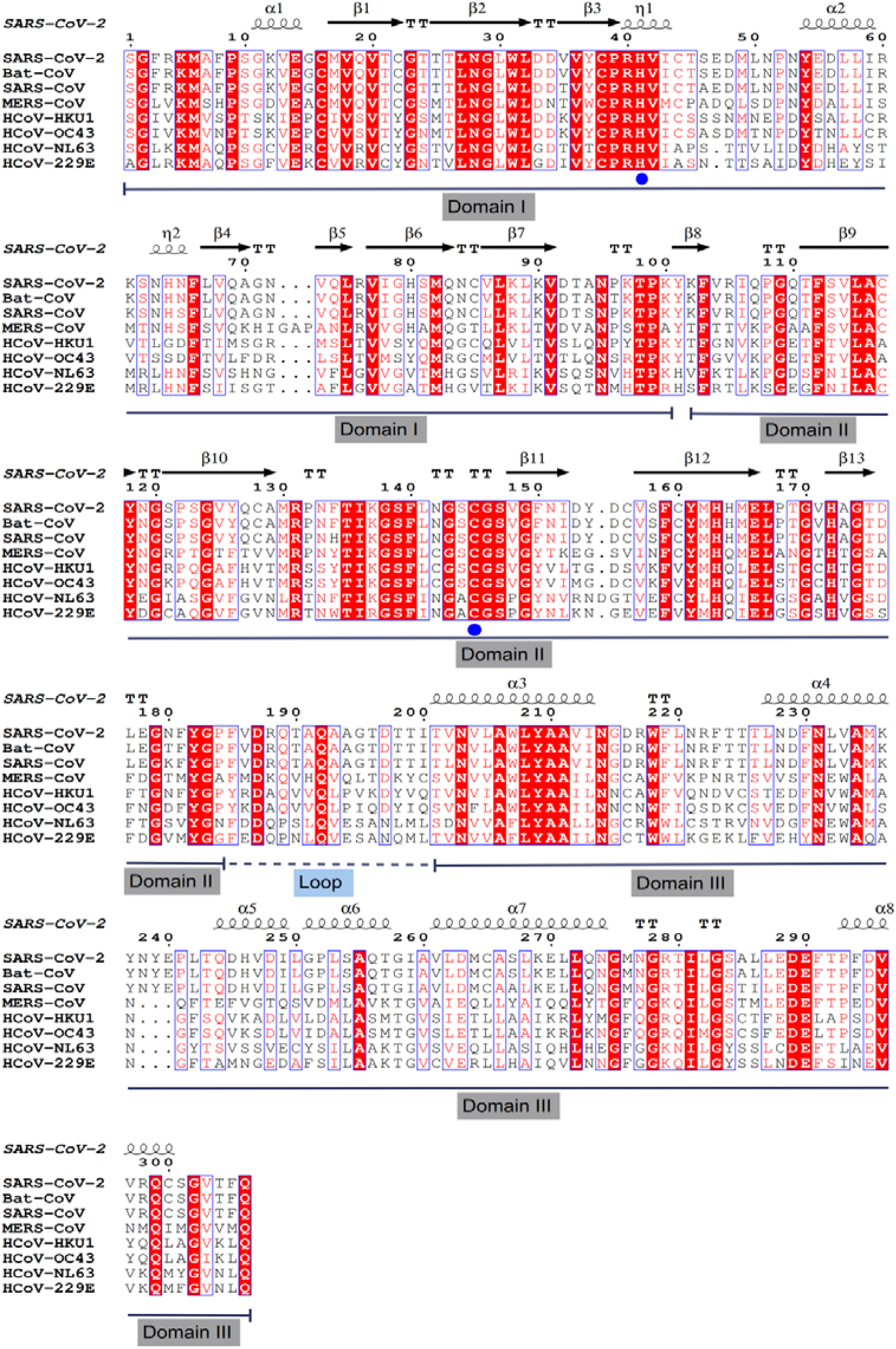
Multiple structure-based sequence alignment of SARS-CoV-2 Mpro was used as a reference and compared with known human coronaviruses: SARS-CoV, Bat-CoV-RaTG13, MERS-CoV, HCoV-HKU1, HCoV-OC43, HCoV-NL63 and HCoV-229E. Domain assignments are marked in the alignment output along with catalytic dyad residues labeled by a blue circle.

**Table 1:** Sequence identity & physiochemical properties of SARS-CoV-2 Mpro with other human coronaviruses Mpro’s.

| Mpro | (%) similarity with SARS-CoV-2 | Length (a.a.) | Mol. Wt. (kDa) | pI | Grand average of hydropathicity (GRAVY) | Instability index |
| --- | --- | --- | --- | --- | --- | --- |
| SARS-CoV-2 | 0 | 306 | 33.8 | 5.95 | -0.019 | 27.65 |
| Bat-CoV-RaTG13 | 99.35% | 306 | 33.8 | 5.95 | -0.007 | 28.36 |
| SARS-CoV | 96.08% | 306 | 33.8 | 6.22 | -0.049 | 29.67 |
| MERS-CoV | 50.65% | 306 | 33.3 | 5.86 | 0.129 | 27.33 |
| HCoV-HKU1 | 49.02% | 303 | 33.2 | 5.78 | 0.16 | 34.51 |
| HCoV-OC43 | 48.37% | 303 | 33.4 | 6.06 | 0.105 | 22.61 |
| HCoV-NL63 | 44.30% | 303 | 32.8 | 6.39 | 0.145 | 31.67 |
| HCoV-229E | 41.04% | 303 | 33.1 | 6.38 | 0.059 | 30.42 |

The majority of (8/12) of variable residues were found in the Mpro β-strands-rich domains I and II, where the inhibitor/catalytic site is located; the remaining (4/12) residues were found in domain III. The connecting loop (residues 185-200) has no variable residues (Figure 1). Also, the SARS-CoV-2 Mpro has only 2 changes when compared to Bat-CoV Mpro and one of the changes is inclusive of 12 changes of Mpro from SARS-CoV and SARS-CoV-2, indicating Bat-CoV as an intermediate between SARS-CoV and SARS-CoV-2 (Figure 2). The finding was consistent with an initial report that SARS-CoV-2 is more similar to SARS-CoV than MERS-CoV, and shares a common ancestor with bat coronaviruses suggesting bat as an intermediate host between SARS-CoV and SARS-CoV-2 [6].

**Figure 2.**
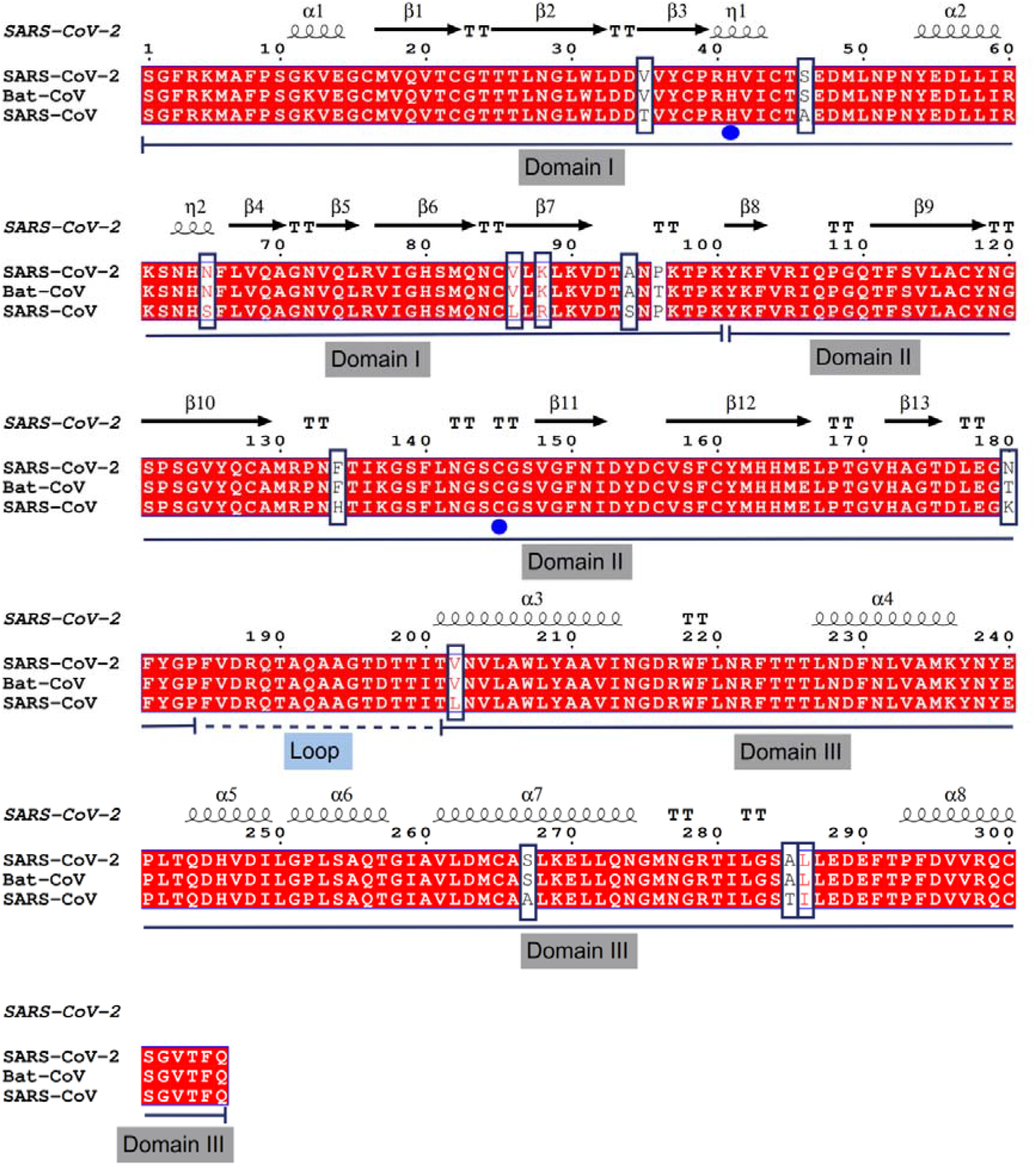
Multiple structure-based sequence alignment of SARS-CoV-2. Mpro was used as a reference and compared with SARS-CoV and Bat-CoV-RaTG13 Mpro. Domain assignments are marked in the alignment output along with catalytic dyad residues labelled by a blue circle. The different residues in the alignment are indicated by the square box.

The 12 different residues in SARS-CoV-2 Mpro compared to SARS-CoV Mpro are V35T, S46A, N65S, V86L, K88R, A94S, F134H, N180K, V202L, S267A, A285T and L286I. Majority of (8/12) of the variable residues were found in the Mpro β-strands-rich domains I and II, the domains where the catalytic site is located; the remaining (4/12) residues were found in domain III. The connecting loop (residues 185-200) has no variable residues (Figure 1). While the substrate binding pockets including the active site residues: T25, T26, H41, M49, F140, N142, G143, S144, C145, H163, H164, M165, E166, P168, H172, Q189, T190, A191 and Q192 are conserved in Mpro from both CoVs. Hence, these above residues will be collectively referred to as active site residues elsewhere in the analysis.

### Structure comparison of Mpro from SARS-CoV and SARS-CoV-2

The available 3D X-ray structure coordinates for SARS-CoV and SARS-CoV-2 Mpro were downloaded from PDB. For our study we used PDB ID: 6M03; (The crystal structure of COVID-19 main protease in an apo form) for SARS-CoV-2, the structure was in apo form i.e. without any other molecule except the protein. While for SARS-CoV Mpro we used PDB ID: 2DUC; (Crystal structure of SARS coronavirus main proteinase (3CLPro)), which was also present in an apo form.

The PDB ID: 2DUC was in dimer form and PDB ID: 6M03 was in monomer form. For comparison of dimer form, the coordinates of 6M03 have transformed accordingly. The SARS-CoV-2 Mpro revealed the presence of three distinct domains: domains I (residues 1–101) and II (residues 102–184) made up by antiparallel β-barrel structures (13 β-strands), connected to the α-helical domain III (residues 201–306) (5 α-helices) by a long loop (residues 185-200) (Figure 3). The catalytic active site is made up of residues from Domain I and II, along with four residues in the long loop region. The dimer interface residues are not involved in active site formation except F140, E166 and H172 which forms key interactions that serve to open and close active site making dimeric form biologically active [32,33].

**Figure 3.**
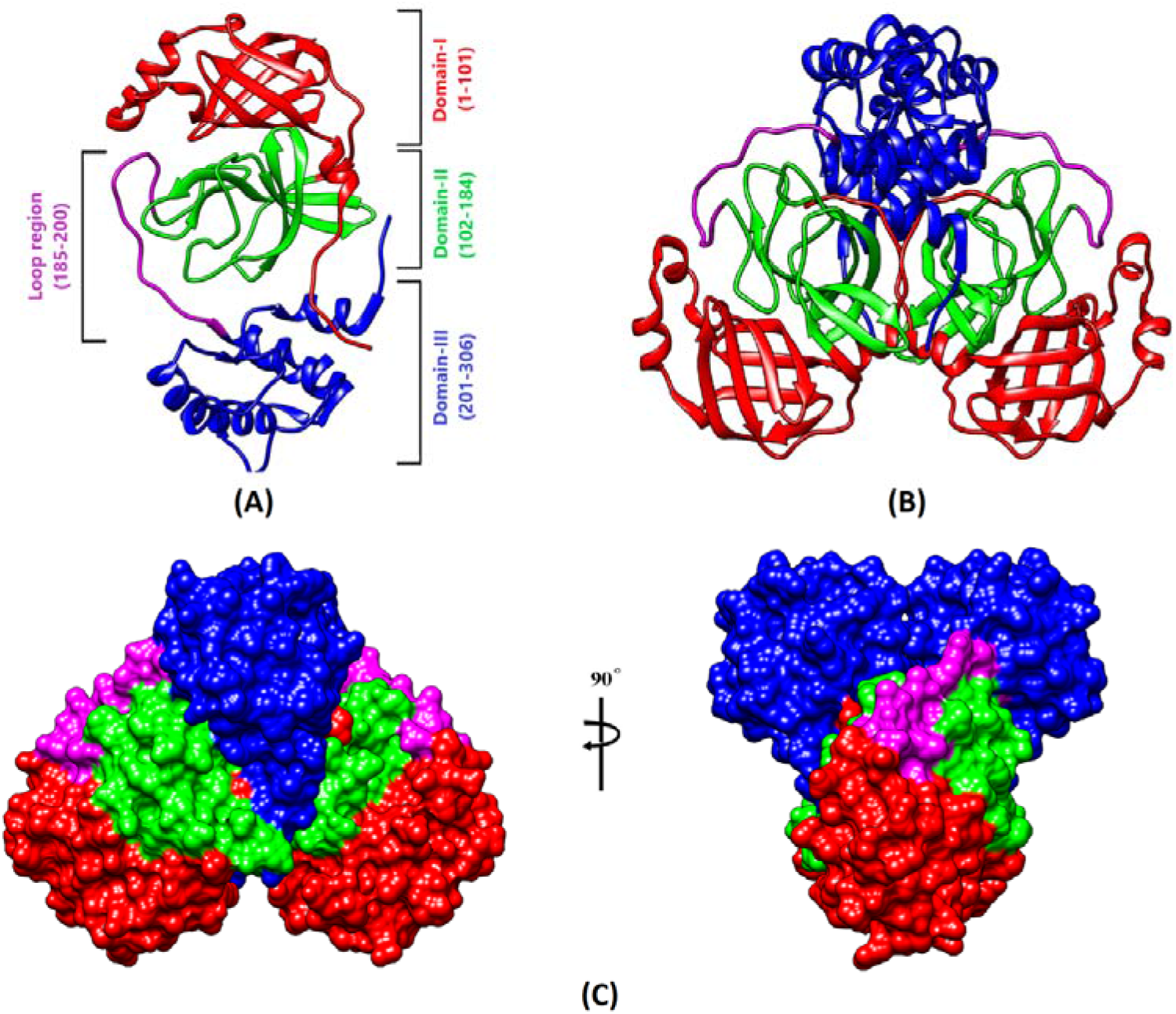
3D structure of SARS-CoV-2 Mpro in monomer and dimer state: (A) Monomer structure of SARS-CoV-2 Mpro with domain assignment marked with residue ranges. (B) Dimer structure of SARS-CoV-2 Mpro coloured by domain assignment. (C) Dimer structure of SARS-CoV-2 Mpro in surface representation coloured by domain assignment at two different angles. In all panels, the Domain-I is marked by red, Domain-II by green, Domain-III by blue and loop region by magenta.

The 3D structure of SARS-CoV-2 Mpro is highly conserved to that of the SARS-CoV Mpro, as expected from the 96% sequence identity; the r.m.s. the deviation is 0.67 Å for all Cα positions (comparison between the two apo-enzyme structures SARS-CoV-2 Mpro; PDB ID: 6M03 and SARS-CoV Mpro; PDB ID: 2DUC). Mpro forms a catalytic active dimer where each monomer consists of the N-terminal catalytic region and a C-terminal region. The dimerization is regulated by Domain III (residues 201-306), a globular cluster of five helices [15,32]. Homo-dimer SARS-CoV-2 Mpro, when superposed on SARS-CoV Mpro, revealed high structural similarity in terms of possessing similar domain orientations and a comparable active site (Figure 4). The tight dimer formed by SARS-CoV-2 Mpro has a contact interface of ~1558 Å and ~1306 Å for SARS-CoV Mpro, between domain II of chain A and the NH2-terminal residues (“N-finger”) of chain B, with the two molecules, oriented perpendicular to one another (Figure 3).

**Figure 4.**
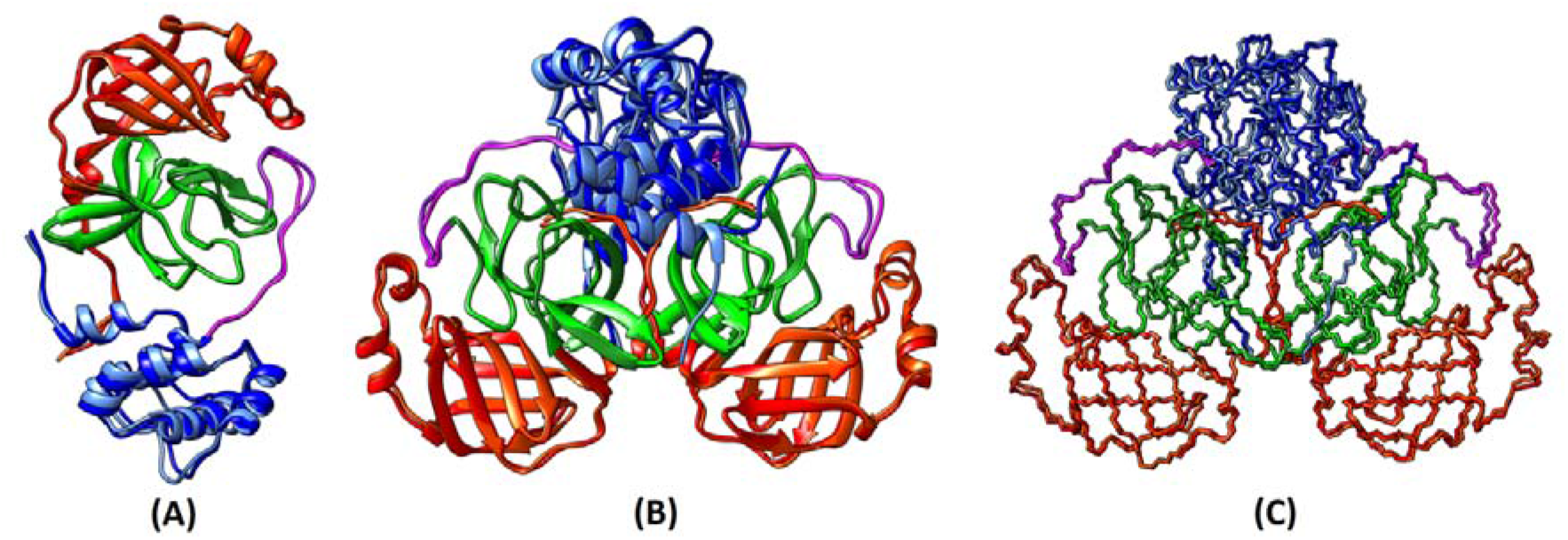
Three-dimensional structure superposition of Mpro from SARS-CoV and SARS-CoV-2 in monomer and dimer state: (A) Monomer structure of SARS-CoV and SARS-CoV-2 Mpro superposed and coloured by domain assignment. (B, C) Dimer structure of SARS-CoV and SARS-CoV-2 Mpro superposed and coloured by domain assignment in cartoon and Cα backbone representation respectively. In all panel, the Domain-I is marked by red (SARS-CoV) & orange (SARS-CoV-2), Domain-II by green (SARS-CoV) & dark green (SARS-CoV-2), Domain-III by blue (SARS-CoV) & light-blue (SARS-CoV-2) and loop region by magenta (SARS-CoV) and purple (SARS-CoV-2).

The 12 different residues (V35T, S46A, N65S, V86L, K88R, A94S, F134H, N180K, V202L, S267A, A285T and L286I) found in Mpro of SARS-CoV-2 compared to SARS-CoV were also analysed for their presence with relative to the active/catalytic site. None of the 12 residues was found in the active site, but only one S46 in SARS-CoV-2 (A46 in SARS-CoV), is located in the proximity of the entrance to the active site (Figure 5). To identify the structural conservancies of SARS-CoV-2 Mpro with other Mpro from alpha, beta, gamma and delta coronaviruses we identify structural homologs available in PDB by BLAST search. The identified Mpro homolog structures from other CoVs are: alphacoronavirus: PEDV-CoV (PDB ID: 5ZQG), HCoV-NL63 (PDB ID: 5GWY), HCoV-229E (PDB ID: 2ZU2), FIPV-CoV (PDB ID: 5EU8), TGEV-CoV (PDB ID: 4F49); betacoronavirus: SARS-CoV (PDB ID: 5B6O), MERS-CoV (PDB ID: 5WKJ), BtCoV-HKU4 (PDB ID: 2YNA), HCoV-HKU1 (PDB ID: 3D23), MHV-A59-CoV (PDB ID: 6JIJ), and gammacoronavirus: IBV-CoV (PDB ID: 2Q6D). Sequence-based structural alignment taking SARS-CoV-2 Mpro as reference structure revealed a striking similarity at the sequence as well as the structural level (Figure 6). All the secondary structural elements for Domain I & II i.e. antiparallel β-barrel structures (13 β-strands), and domain III i.e. 5 α-helices were found to be present in all CoV Mpro (Figure 6). Phylogenetic analysis of these 12 sequences as well as newly identified BatCoV-RaTG13 Mpro revealed clear distinct clades made up of alpha, beta and gamma coronavirus. The Mpro from SARS-CoV and SARS-CoV-2 form a distinct clade different from rest of the sequences and the latter one being very close to recently identified BatCoV-RaTG13 Mpro [6].

**Figure 5.**
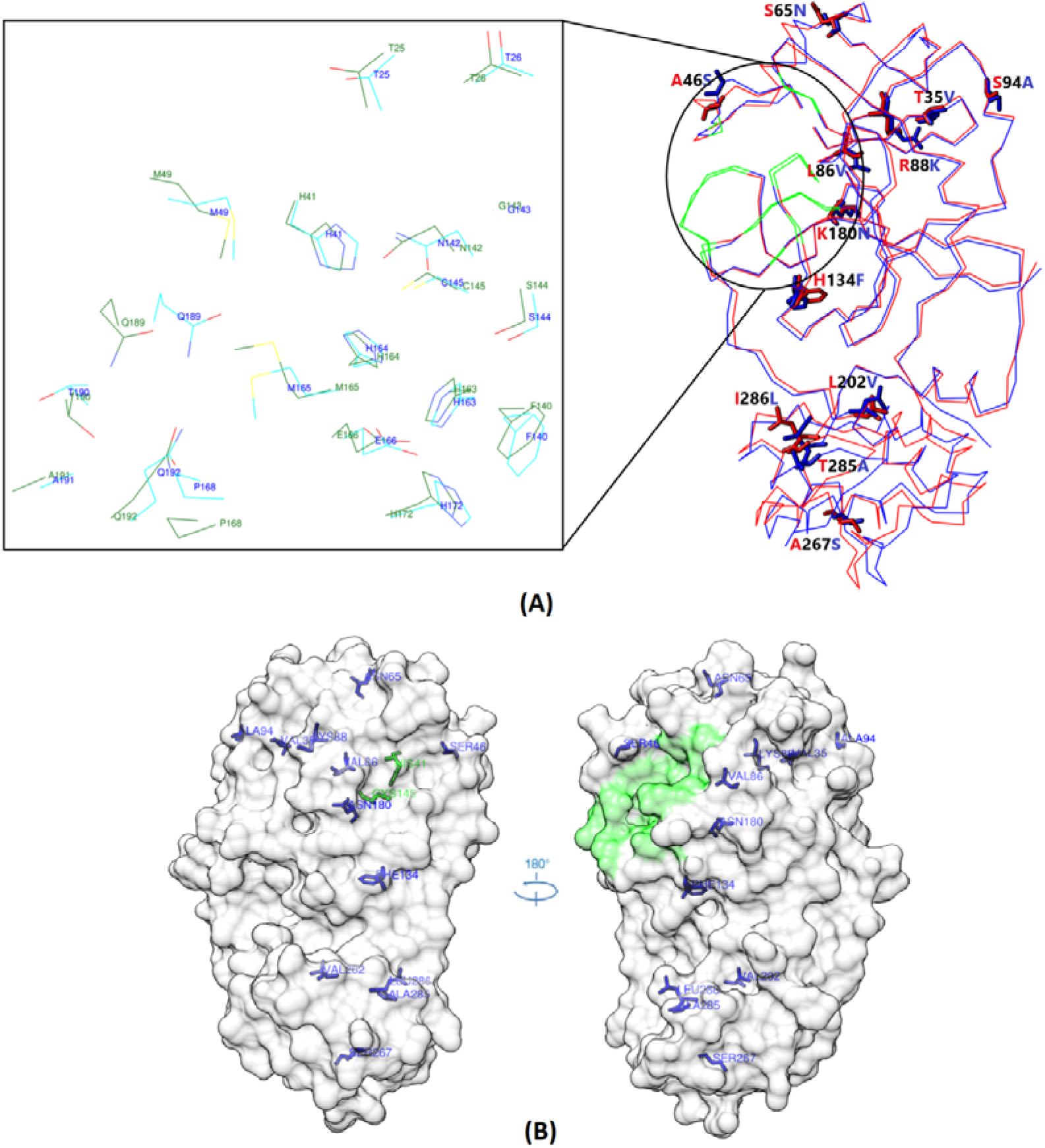
Monomer structure alignment of SARS-CoV Mpro (red) and SARS-CoV-2 Mpro (blue) aligned based on Cα backbone. (A) The overall structure of both SARS-CoV and SARS-CoV-2 Mpro with 12 differing amino acids marked and shown in sticks as red (SARS-CoV Mpro) and blue (SARS-CoV-2 Mpro). The active sites residues are coloured in green. Close-up of active site residues and represented in sticks (side chains) as cyan (SARS-CoV-2 Mpro) and dark green (SARS-CoV Mpro). (B) Structure of SARS-CoV-2 Mpro in surface display in two angles (180° rotation) with 12 different residues from SARS-CoV Mpro marked and shown in blue sticks. The catalytic HIS41 and CYS145 residues are shown in green sticks and other residues of the active site are shown on the green surface.

**Figure 6.**
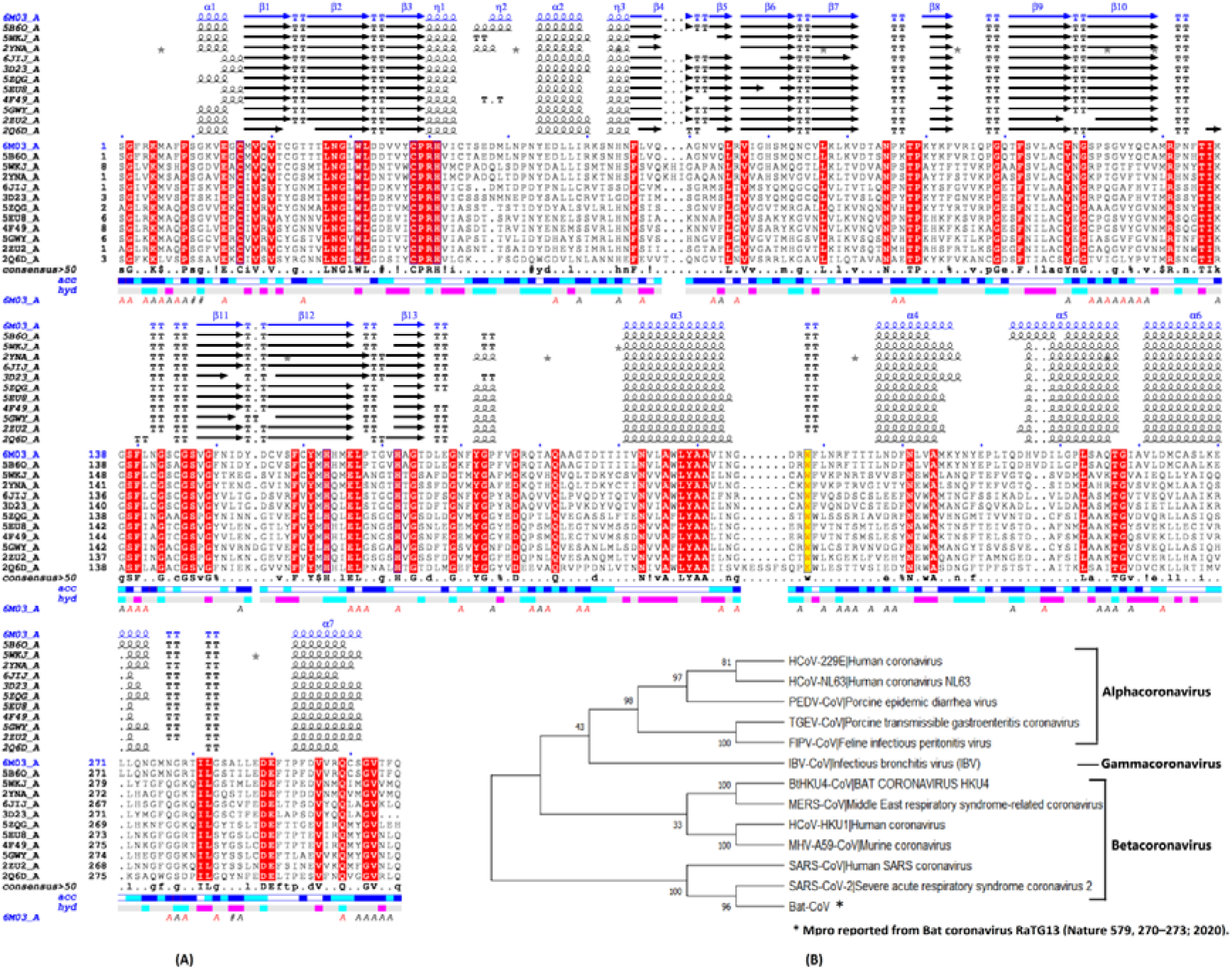
Multiple Sequence Alignment and phylogenetic analysis of Mpro from 12 different CoV: (A) Multiple structure-based sequence alignment of SARS-CoV-2 Mpro taking as a reference and its comparison with known coronaviruses: SARS-CoV, BAT-CoV-KHU4, MERS-CoV, HCoV-HKU1, HCoV-NL63, HCoV-229E, MHV-A59-CoV, PEDV-CoV, TGEV-CoV, FIPV-CoV, IBV-CoV and BatCoV-RaTG13. (B) Phylogenetic tree construction by Maximum Parsimony method using above 12 Mpro sequences as well as a recently reported Mpro sequence from BatCoV RaTG13.

Looking at the RMSD (Å) of the superposed 12 Mpro crystal structures, it was suggested that most flexible regions was helical domain III (residues 201–306) and loops on the surface. However, substrate-binding pockets were located in a cleft between domains I and II, which were still highly conserved among all CoV Mpros, suggesting the feasibility of designing antiviral inhibitors targeting this site (Figure 7). It shows a variable tube representation of the Cα trace of 12 Mpro homologous protein structures superposed onto the SARS-CoV-2 Mpro; PDB ID: 6M03 and the size of the tube is proportional to the mean r.m.s. deviation per residue between Cα pairs. The blue to red colour ramping shows sequence conservation from high to low. The analysis revealed identifies areas of weak and strong structural conservation correlating with the sequence similarity in substrate-binding pockets located in a cleft between domains I and II (Figure 7). Contrary, on sequence level variation (8/12) residues are found in Mpro β-strands-rich domains I and II, where the inhibitor/catalytic site is located; the remaining (4/12) residues were found in domain III. Our analysis indicates that though the variations found in SARS-CoV-2 Mpro compared to SARS-CoV didn’t confer flexibility to the region where the inhibitor/catalytic site is located based on RMSD (Å) deviation of Cα atoms.

**Figure 7.**
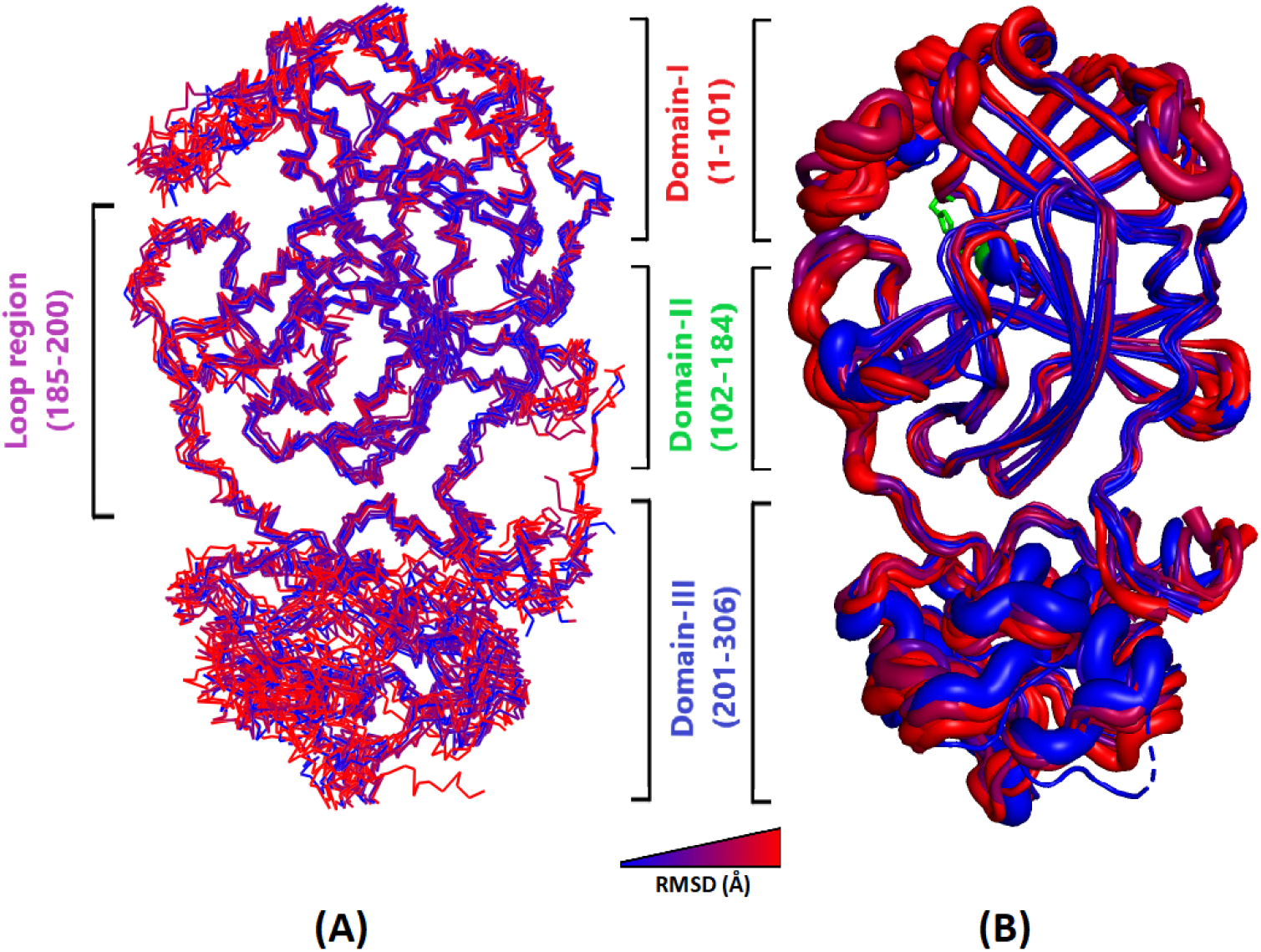
Structural superposition of 12 CoV Mpro from different strains. (A) Superposition of 12 Mpro structures (Cα backbone) from different CoV onto SARS-CoV-2 Mpro as a query. (B) Representation of 12 Mpro superposed structures using variable tube depiction, where the radius is proportional to the RMSD differences in Cα between SARS-CoV-2 Mpro and 12 other homologous Mpro structures. In both the panel, blue to red colour ramping is used to visualize and correlate conservation from areas of strong to weak conservation respectively. Domain assignments are marked with residue ranges in both the panels.

### Molecular Dynamics Simulations (MDS) of Mpro from SARS-CoV-2 and SARS-CoV

The simulations were carried out to study the conformational behaviour, structural dynamics, and stability of the Mpro of SARS-CoV-2 and SARS-CoV. We analysed the RMSD, RMSF, H-bonds, as well as PCA properties of both the dimeric and monomeric forms. A series of gmx commands and analysis were run for all four (SARS-CoV_ monomer, SARS-CoV_dimer, SARS-CoV-2_monomer and SARS-CoV-2_dimer) MDS to interpret the difference between Mpro from SARS-CoV-2 and SARS-CoV for single configurations according to functions like RMSD, RMSF, ROG etc. by obtaining value/s for each time point throughout the trajectory. Subsequently, dynamics of both Mpro were also investigated in the time domain by averaging the fluctuations across entire simulations.

### Stability of Mpro from SARS-CoV-2 and SARS-CoV

The overall stability of Mpro was investigated through MDS run for assessing configurational interpretation at each time-point. First RMSD values for the entire 50 ns simulations were evaluated to access the convergence of the simulation towards equilibrium in terms of the Euclidean distance from the average structure to a reference (crystal) structure. It is evident from the plots shown in Figure 8, that most of the MDS have reached equilibrium after 25 ns. The mean RMSD values of protein Cα backbones for SARS-CoV_monomer, SARS-CoV-2_monomer, SARS-CoV_dimer and SARS-CoV-2_dimer were 0.17 ± 0.035, 0.17 ± 0.025, 0.17 ± 0.020 and 0.13 ± 0.013 nm respectively. The mean RMSD values for Mpro monomer for both the CoV as well as Mpro from CoV dimer have similar values. The lower RMSD values of SARS-CoV-2_dimer compared to Mpro monomer as well as SARS-CoV_dimer indicated the formation of a stable molecule. Further, all four MDS were subjected to Radius of Gyration (ROG) calculation for the entire 50 ns simulations. ROG is a measure of shape of the protein at each time-point by comparing it to the experimentally obtainable hydrodynamic radius. The mean ROG values of protein Cα backbones for SARS-CoV_monomer, SARS-CoV-2_monomer, SARS-CoV_dimer, and SARS-CoV-2_dimer were 2.18 ± 0.011, 2.18 ± 0.014, 2.55 ± 0.0081 and 2.54 ± 0.0074 nm, respectively (Figure 8). The ROG values of Mpro were similar w.r.t to monomer and dimer forms in both the CoVs indicating that the overall shape of a protein is consistent irrespective of the changes at the residue level.

**Figure 8.**
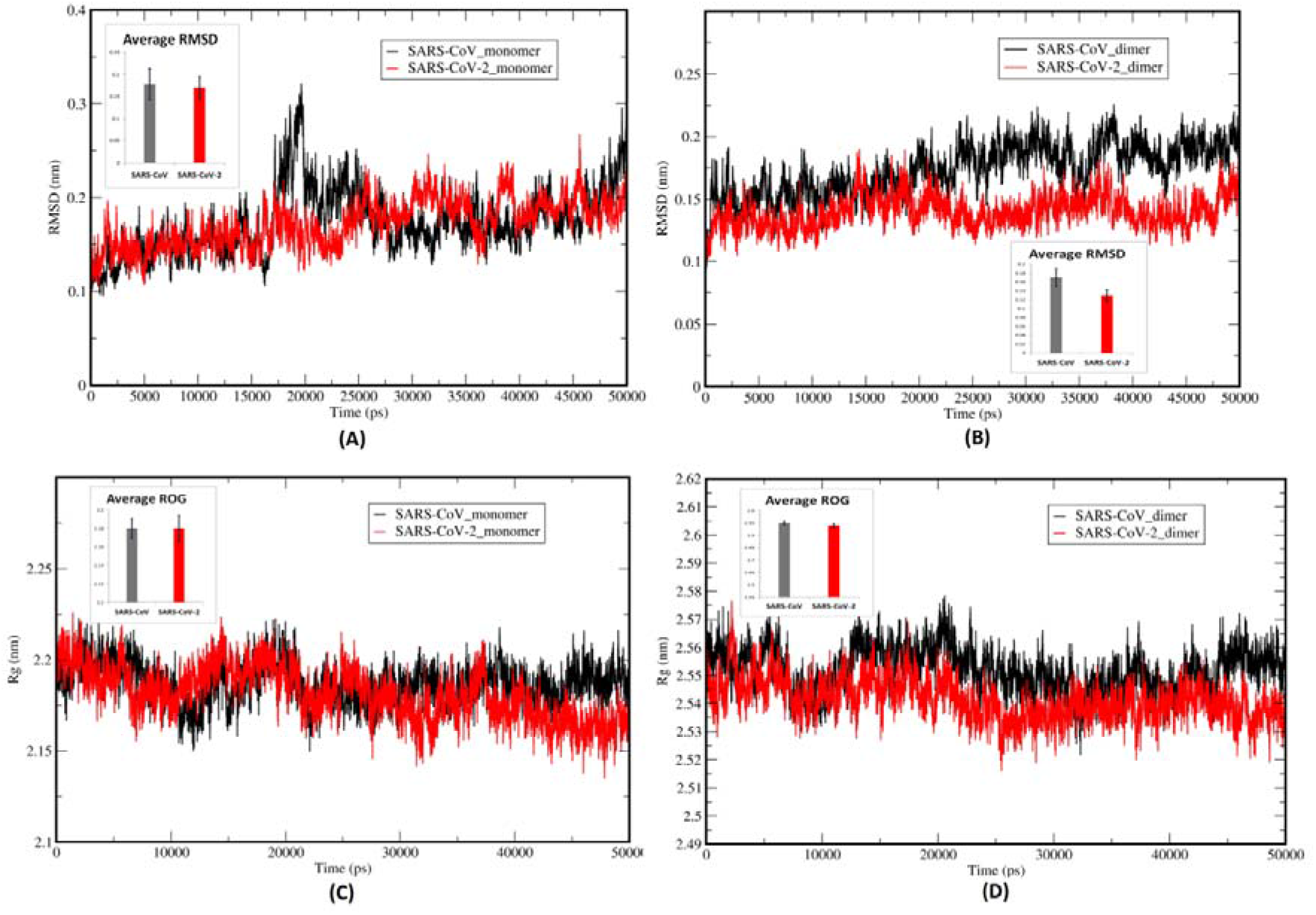
Molecular Dynamics Simulations of Mpro from SARS-CoV and SARS-CoV-2 in monomer and dimer structures, computing the deviation (nm) vs. function of time (50 ns): RMSD of the protein Cα backbone atoms of SARS-CoV and SARS-CoV-2 Mpro in monomer (A) and dimer (B) forms. ROG of the protein Cα backbone atoms of SARS-CoV Mpro and SARS-CoV-2 Mpro in monomer (C) and dimer (D) forms. Inset graph represents average values with standard deviations and the values of SARS-CoV and SARS-CoV-2 Mpro are plotted in black and red colour respectively.

We also reported the formation of H-bonds for entire simulation trajectories for MD systems. The secondary structures which form the core of protein structures underpinning protein fold are stabilized by H-bonds and thus are an indicator of rigidity to the protein structure and specificity to intermolecular interactions between secondary structures. The mean H-bond values for intra-molecular interactions for SARS-CoV_monomer, SARS-CoV-2_monomer, SARS-CoV_dimer, and SARS-CoV-2_dimer were 211.73 ± 6.96, 217.415 ± 7.25, 441.363 ± 9.83 and 451.954 ± 9.92 (Figure 9). The H-bond analysis indicated no substantial difference between Mpro from either CoV in both the dimer and monomer form. To access the surface area of protein that is accessible to solvent in which it is simulated, we calculated the surface accessible solvent area (SASA) variable for the entire trajectory. Total SASA calculated for SARS-CoV_monomer, SARS-CoV-2_monomer, SARS-CoV_dimer and SARS-CoV-2_dimer were 150.48 ± 2.28, 149.38 ± 2.18, 272.744 ± 3.18 and 265.811 ± 2.65 (Figure 9). For Mpro monomer there is no significant difference between SARS-CoV and SARS-CoV-2, but for the dimer, the SARS-CoV-2 Mpro had little lesser SASA values compared to SARS-CoV indicating a lesser magnitude of flexibility and instability.

**Figure 9.**
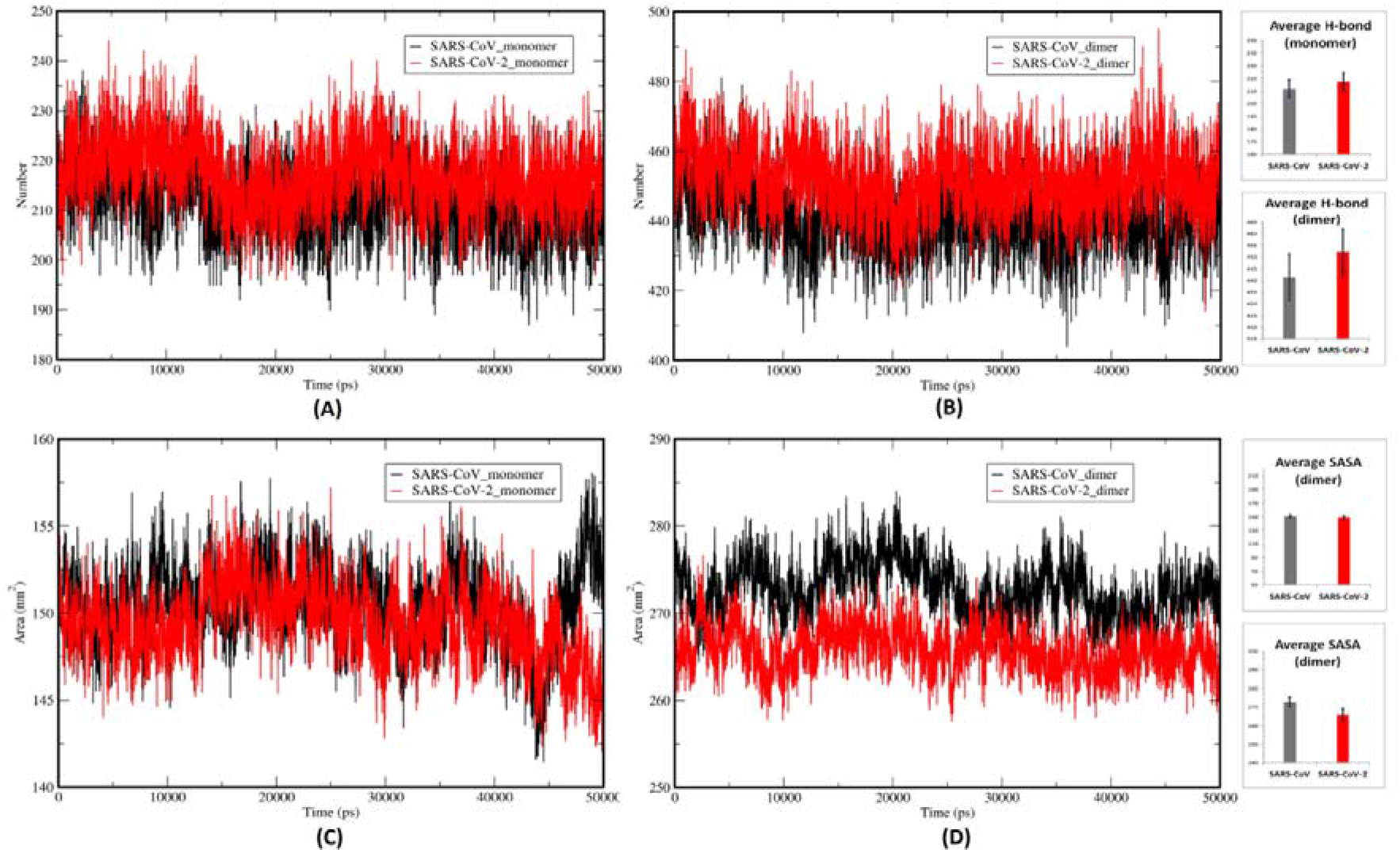
Intra hydrogen bonds and Solvent Accessible Surface Area (SASA) of Mpro from SARS-CoV and SARS-CoV-2 Mpro in monomer and dimer structures: Number of hydrogen bonds computed vs. function of time (50ns) in Mpro of SARS-CoV and SARS-CoV-2 in monomer (A) and dimer (B) forms. Total SASA values plotted (nm) for entire simulations in Mpro of SARS-CoV and SARS-CoV-2 in monomer (C) and dimer (D) forms. Inset graph represents average values with standard deviations and the values of SARS-CoV and SARS-CoV-2 Mpro are plotted in black and red colour respectively.

The SASA values for 12 different residues as well as the active site residues are plotted in Figure 10 for all the four MD systems. For residues V35T, N65S AND F134H the solvent-exposed surface area was higher in SARS-CoV-2 compared to SARS-CoV. The variable residue N65S is close to the binding site (T25, T26, M49 and Q189). Variable residue F134H is also critical with the fact that many functionally crucial residues [H172, E166, F140 (residues also involved in dimerization) and the oxyanion loop (140-144)] are found in its vicinity. F134H is also in the loop that leads to the catalytic residue C145. Alternatively, for residues K88R, N180K and V202L, the SASA values were lower in SARS-CoV-2 compared to SARS-CoV. The variable residue K88R and N180K are close to the catalytic site.

**Figure 10.**
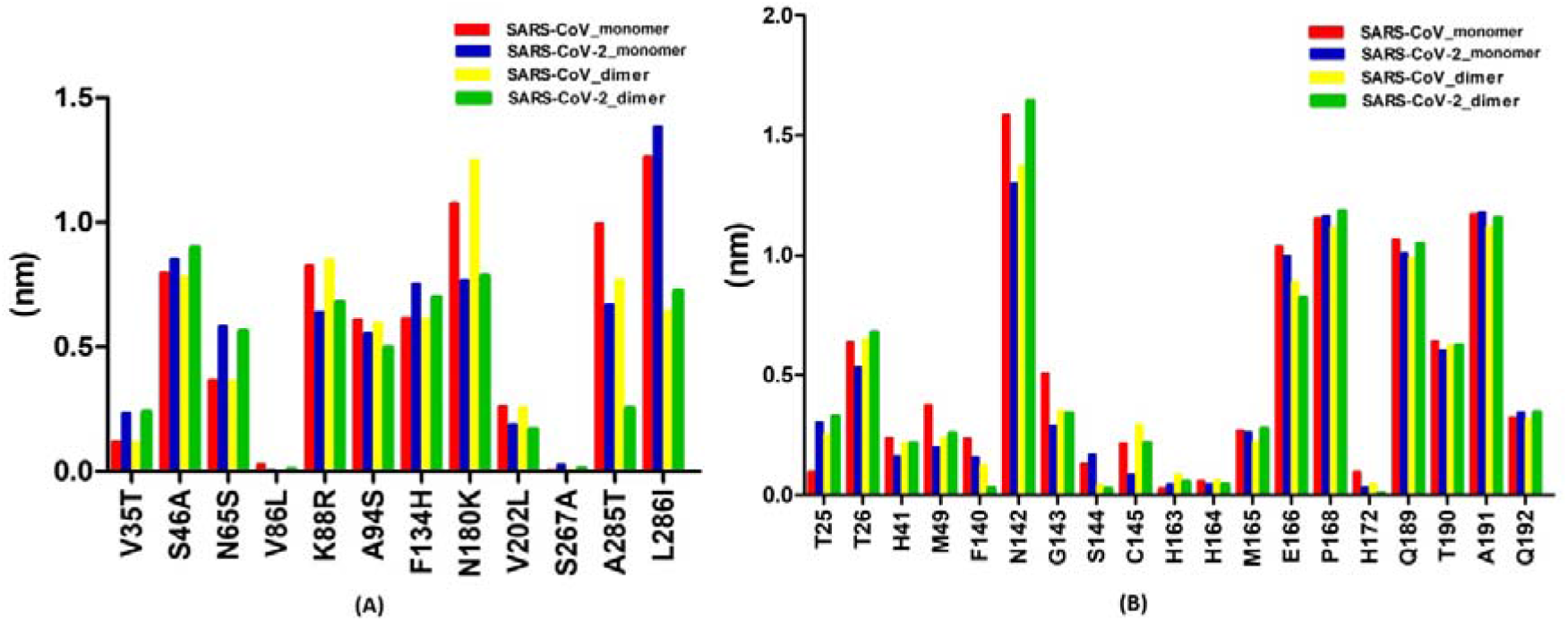
The plot of SASA values for individual residues in all four MD systems: (A) The plot of 12 divergent residues in SARS-CoV-2 Mpro. The X-axis represents the residue present in SARS-CoV-2 followed by the residue number and ending with the residue found in SARS-CoV. (B) The plot of residues present in active site which are conserved in SARS-CoV and SARS-CoV-2 Mpro.

The SASA values for A285T and L286I were very different and lesser for dimer when compared to its monomer counterpart. Specifically, the values for A285T were very lower in case of SARS-CoV-2_dimer compared to SARS-CoV_dimer and this is correlated with the reduction of the distance between Cα of A285 in SARS-CoV-2_dimer which allows the two domains III to approach each other a little closer, also supported by an earlier finding [16,33]. Our findings suggest that despite having overall structural similarity with the SARS-CoV Mpro, the 12 divergent residues in SARS-CoV-2 Mpro might affect the catalytic activity by modifying the microenvironment of the catalytic site, conferring a potential change in SARS-CoV-2.

The SASA values for active site residues were also compared to compute the change in their exposure to solvent vis. a vis. 12 different residues. The values for T25 for dimer were larger than monomer in both the Mpro. While for M49 and G143, the values for SARS-CoV2 monomer as well dimer were lesser than SARS-CoV Mpro. T25 and M49 are very close to the variable position 46 and 65 and possibly affected by them. The residues at F140, H172 and E166 in case of SARS-CoV2 dimer had very lesser values compared to SARS-CoV dimer and these key residues are also involved in dimerization and vicinity to a variable residue F134H in SARS-CoV-2 Mpro. The catalytic residue at C145 had lower SASA value in SARS-CoV2 than SARS-CoV in both the monomer as well dimer form. Notably variable position S46A and F134H are found in a loop that leads to this catalytic H41 and C145 respectively.

### Residual Fluctuations and molecular interactions in Mpro from SARS-CoV and SARS-CoV-2

The Root Mean Square Fluctuations (RMSF) values are an important measure for each atom’s fluctuation about its average position. RMSF analysis reveals important insight into the flexibility of regions of the molecule. RMSF plots for each MD systems are shown in (Figure 11). The calculated average RMSF values for SARS-CoV_monomer, SARS-CoV-2_monomer, SARS-CoV_dimer and SARS-CoV-2_dimer were 0.14 ± 0.054, 0.10 ± 0.056, 0.1015 ± 0.037 and 0.0825 ± 0.029 nm respectively. The SARS-CoV-2 Mpro had overall lesser RMSF values and particularly the SARS-CoV-2_dimer having the least indicating higher stability and lesser fluctuations. The RMSF analysis of protein backbone in CoV_monomer revealed that the domain I followed by domain II region to be one of the highly flexible regions in case of SARS-CoV_monomer. As shown in figure 11, fewer fluctuations in domain I and II in case of SARS-CoV-2_monomer is observed, indicating the formation of a more stable molecule. In the case of SARS-CoV-2_monomer, a continuous stretch of residues from 20-110 is found to have very fewer fluctuations compared to SARS-CoV_monomer. Unlike, the SARS-CoV-2_monomer, the SARS-CoV-2_dimer overall RMSF values for each domain was less compared to SARS-CoV_dimer.

**Figure 11.**
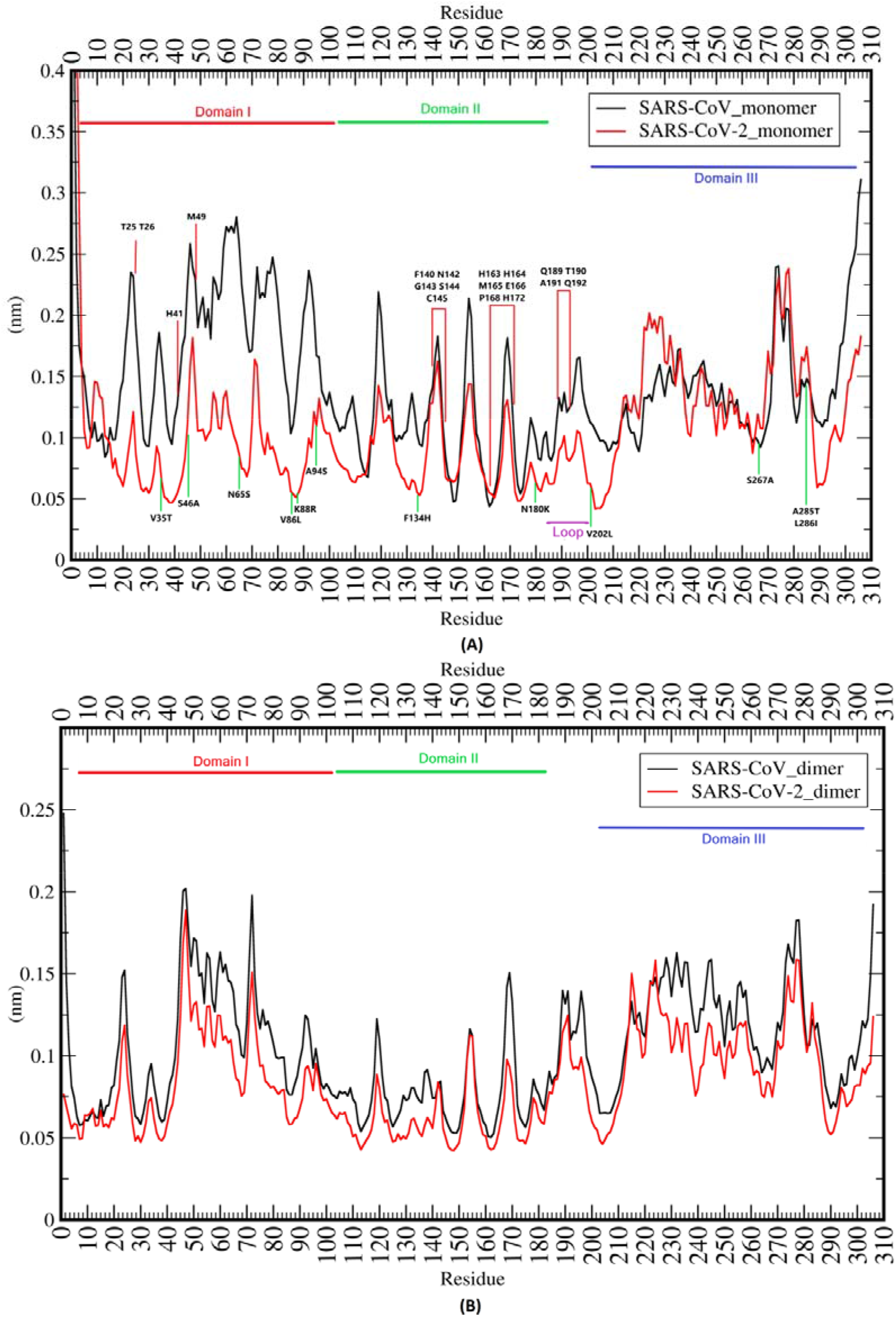
Residue-wise RMSF deviations (nm) of Mpro from SARS-CoV and SARS-CoV-2 in monomer and dimer form: (A) RMSF plot of both CoV Mpro in monomer form. The 12 divergent residues found in SARS-CoV-2 are marked with a green line in the lower plot. The active site residues are marked with a red line in the upper plot. (B) RMSF plot of both CoV Mpro in dimer form. In both the panels, the domain I, II and III residues are marked in red, green and blue colour respectively.

Further, we analysed the individual RMSF values of the 12 divergent residues and the residues forming active site in Mpro and plotted as shown in Figure 12. In the case of SARS-CoV-2_monomer, all the 12 divergent residues showed a significant decrease in fluctuations (RMSF) than SARS-CoV_monomer Mpro, except the three variant S267A, A285T and L286I which are found in domain III. Alternatively, in both CoV dimer, all the variant residues in SARS-CoV-2 showed fewer fluctuations compared to SARS-CoV. The above three variants present in domain III also showed lesser values, maybe due to dimerization implications. Notably, in case of SARS-CoV-2_monomer Mpro, the V35T, N65S, K88R, F134H and V202L residues had at least 50% lesser RMSF values compared to SARS-CoV_monomer. The variable residue N65S is close to the binding site (T25, T26, M49, and Q189); K88R is close to the catalytic site; F134H is present in the loop that leads to catalytic C145 as well as H172, E166, F140 (residues involved in dimerization) are present in its vicinity.

**Figure 12.**
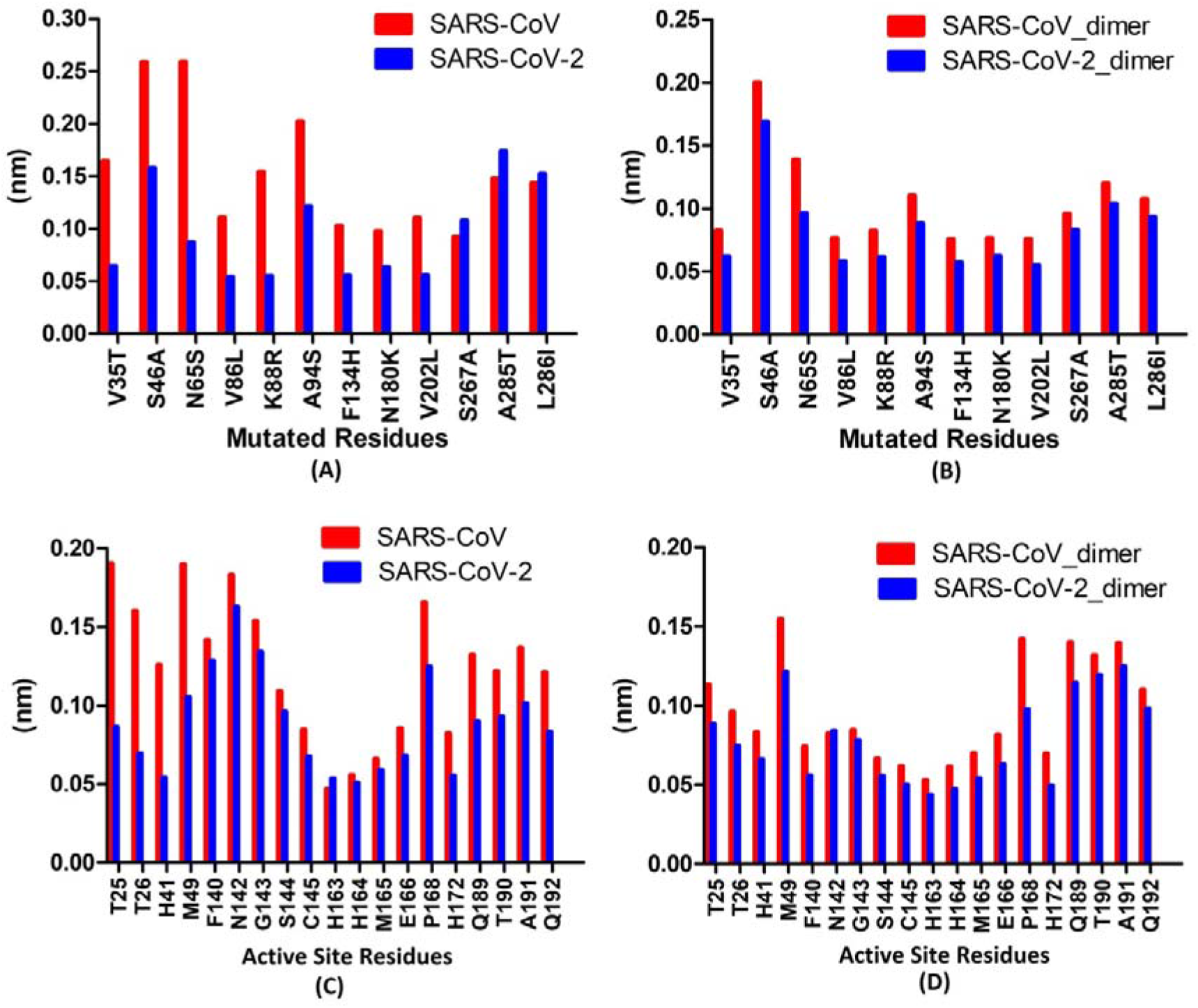
RMSF plots showing deviations (nm) of selected residues from SARS-CoV and SARS-CoV-2 Mpro in monomer and dimer form: RMSF plots of 12 divergent residues from both the CoV Mpro in monomer (A) and dimer (B). The X-axis represents the residue present in SARS-CoV-2 followed by the residue number and ending with the residue found in SARS-CoV. RMSF plots of active site residues from both the CoV Mpro in monomer (C) and dimer (D).

The individual RMSF values of residues from the active site also showed a similar trend as observed in the divergent 12 residues (Figure 12). In the case of monomer Mpro, the residues of domain I showed significantly lesser values in SARS-CoV-2_monomer as compared to SARS-CoV_monomer. In the case of SARS-CoV-2_monomer Mpro, the T25, T26, H41 and M49 residues had significantly lesser RMSF values compared to SARS-CoV_monomer. Notably, these environments of the above residues are directly affected by the variable residues like K88R, N65S, and S46A. Unlike the monomer CoV, in case of both the CoV dimer, all the active site residues in SARS-CoV-2_dimer showed fewer fluctuations compared to SARS-CoV_dimer. A similar trend is also observed in case of RMSF values of the 12 variant residues. Surprisingly, the residues involved in dimerization (F140, H172, and E166) didn’t show significant differences in RMSF in both the monomer and dimer CoV. Overall, the results of RMSF analyses were in correlation with the SASA analysis.

Molecular interactions for key residues in both the CoV Mpro (monomer and dimer) were analysed using foldx program from the final structure obtained after MD simulations. We assess the change in molecular interactions or network in key residues which may have been introduced due to mutation in SARS-COV-2. The variable position at S46A is located near the entrance of the binding site (Figure 5) has lost its interaction with M49 (active site residue) in both the monomer and dimer form of SARS-CoV-2 Mpro as also evident from earlier results [34]. The variable residue at position F134H results in a substitution of +ve charge H134 to a hydrophobic F134 that altered the environment, and the fact that it is located at the loop leading to the oxyanion loop, it may serve to modulate active site (unique interaction to SARS-CoV-2: A105 and SARS-CoV: G183). Variable residue at V86L which is located near the catalytic site has resulted in a loss of interaction in SARS-CoV-2 (SARS-CoV: G179). Another variable position K88R located near the catalytic site has altered the electrostatic interaction profile in SARS-CoV-2 (unique to SARS-CoV-2: E55 & H164 and SARS-CoV: F103 & K180). The variable A94S position has resulted in a loss of its interaction with P96 in SARS-CoV-2. Interestingly, the variable position at N180K has resulted in a loss of all electrostatic interactions in SARS-CoV-2 due to the mutation of +ve charged residue (K) to a polar uncharged residue (N). Otherwise K180 in SARS-CoV- had electrostatic interactions with R40, R88, R105, D176, E178, and D187. The variable residues A285T and L286I in SARS-CoV-2 has resulted in a change in interaction as compared to SARS-CoV as well as make new connections with T280 (chain B) and G283 (chain B), at the interface of the dimer.

The catalytic residues H41 and C145 didn’t have a much-altered environment due to the mutations found in SARS-CoV-2. However, a new network of H-bond was formed in case of SARS-CoV-2: H41 with H164, which is also one of the residues involved in active site formation. Such N-H···N hydrogen bonds formed by imidazole groups of two histidine residues (H41: H164) are rare in proteins [35]. Our SASA analysis indicated that H41 & H164 are relatively buried and such N-H···N H-bonds formed by a pair of buried histidine, may significantly contribute to structural stability [35] of the SARS-CoV-2 Mpro which is evident by our above RMSD and RMSF analysis. There is no change in the network of C145 catalytic residues which indicates that the integrity of C145 may be very essential for its conserved protease activity. The M49 and Q189 present at the entrance of the active site are essential gatekeepers for substrate binding [36]. The network of M49 has been changed slightly in SARS-CoV-2 Mpro (gained in SARS-CoV-2: T45 & Q189 and lost: P52 & D187). Similarly in the other gatekeeper residue Q192, its interaction with T190 was lost in SARS-CoV-2 while interaction with M165 was found unique to SARS-CoV-2. A new interaction was gained by G143 to N28 in case of SARS-CoV-2 which is absent in SARS-CoV. Similarly, a new interaction was gained by H163 to M165 in case of SARS-CoV-2 which is absent in SARS-CoV. An important residue, M165 found adjacent to the key residue E166 which is required to open the substrate-binding site in CoV Mpro has gained a unique connection in SARS-CoV-2 (HISA163, VALA186, ASPA187 and GLNA192) which are completely absent in SARS-CoV Mpro. A new interaction was gained by P168 to T169 in case of SARS-CoV-2 which is absent in SARS-CoV. For position T190, a potential network with Q192, another active site residue was lost in case of SARS-CoV-2 Mpro.

The Mpro dimer form is reported to be functional due to crucial interactions of the residues F140, E166, H172 and H163 within and between the residues of another chain specifically the N-finger which serve to open and close the active site as well as an active role in dimerization [15,32]. We analysed the interaction network of this important residue involved in dimerization. Our analysis revealed that the key interaction of F140, E166 and E290 with N-finger residues of other chain is maintained in particular with residues S1 and R4 of another chain. In case of E166, apart from the N-finger residues, it also interacts with N214, D216 and C300 of another chain in both SARS-CoV and SARS-CoV-2 Mpro but a new unique interaction to SARS-CoV-2: R217 is also observed. Interestingly, S1 from N-finger of other chain forms a unique H-bond with H172 in SARS-CoV-2 but absent in SARS-CoV. H172, as it is one of the key residues in the active site of CoV Mpro which makes a typical H-bond with S1 of other chain, may also contribute to the restructuring process of SARS-CoV-2 Mpro. E290 is an also crucial residue which makes an interaction with R4 of other chain forming a salt bridge interaction and the same was also observed in both the CoV Mpro. The collective analysis by RMSF, SASA and molecular interactions revealed that the 12 divergent residues in SARS-CoV-2 Mpro alter the micro-environment of neighbouring residues. These modified interaction networks finally restructure the molecular environment of the Mpro active site residues at the entrance (T26, M49, and Q192) and near the catalytic region (F140, H163, H164, M165 and H172).

### Clustering of conformations for ensemble generation and Essential Dynamics

The conformational space and transitions in the SARS-CoV and SARS-CoV-2 Mpro for monomer and dimer were inspected by Principle Component Analysis (PCA) analysis. The PCA is a statistical computation that decreases the complexity of the MDS trajectories by extracting only the collective motion of Cα atoms while preserving most of the other variation. It calculates the covariance matrix of positional fluctuations for backbone atoms which may decipher the dynamics and coherted motions of Mpro from both SARS-CoV and SARS-CoV-2. Figure 13 showed a plot of eigenvalues calculated from the covariance matrix of backbone fluctuations, plotted in decreasing order vs. the respective eigenvector indices for all MD systems. Top 15 eigenvectors accounted for 77.50 %, 75.36 %, 68.08 % and 60.83 % of motions observed for 50 ns trajectory for SARS-CoV_monomer, SARS-CoV-2_monomer, SARS-CoV_dimer, and SARS-CoV-2_dimer respectively. The plot which is shown in Figure 13 is the 2D projection of the trajectories for two major principal components PC1 and PC3 for SARS-CoV_monomer, SARS-CoV-2_monomer, SARS-CoV_dimer, and SARS-CoV-2_dimer which represents different conformations in 2D space. The PCA analysis revealed the following observations. First, the 2D projection of SARS-CoV_monomer (Figure 13B) has more variation compared to the other three: SARS-CoV-2_monomer (Figure 13C), SARS-CoV_dimer (Figure 13D), and SARS-CoV-2_dimer (Figure 13E). Second, it is evident from the 2D plot that the SARS-CoV-2_monomer (Figure 13C) and SARS-CoV-2_dimer (Figure 13E) showed higher stability and occupies lesser phase space compare to SARS-CoV_monomer (Figure 13B) and SARS-CoV_dimer (Figure 13D). This indicates that Mpro of SARS-CoV-2 in stable in both the form and less flexible than SARS-CoV Mpro. The covariance and 2D plot analysis also indicated the presence of two well-defined clusters in SARS-CoV_monomer, SARS-CoV-2_monomer & SARS-CoV_dimer and only one defined cluster in case of SARS-CoV-2_dimer. The positive and negative limits are depicted by the covariance plots where; positive values are related to the motion of the atoms occurring along the same direction (correlated), whereas the negative values indicate motion of the atoms in the opposite direction (anti-correlated). Our PCA analysis from MD simulations (50 ns) revealed that the SARS-CoV_monomer (Figure 13B) and SARS-CoV_dimer (Figure 13D) had more anti-correlated motion and the SARS-CoV-2_monomer (Figure 13C) and SARS-CoV-2_dimer (Figure 13E) had a balance of correlated as well as anti-correlated motion. Thus, from the above results, it was concluded that SARS-CoV_monomer and SARS-CoV_dimer had increased flexibility and conformational space compared to the SARS-CoV-2_monomer and SARS-CoV-2_dimer and notably, the most stable and less flexible was of SARS-CoV-2_dimer. To visualize the essential dynamics, 50 structures were extracted from each MD simulation projecting the extremely selected eigenvectors (Figure S1). The extreme motion of SARS-CoV-2_monomer and SARS-CoV-2_dimer were less deviating when compared to SARS-CoV_monomer and SARS-CoV_dimer, indicating a stable conformational space. The width of the main-chain trace represents fluctuations throughout the timescale of MD simulations.

**Figure 13.**
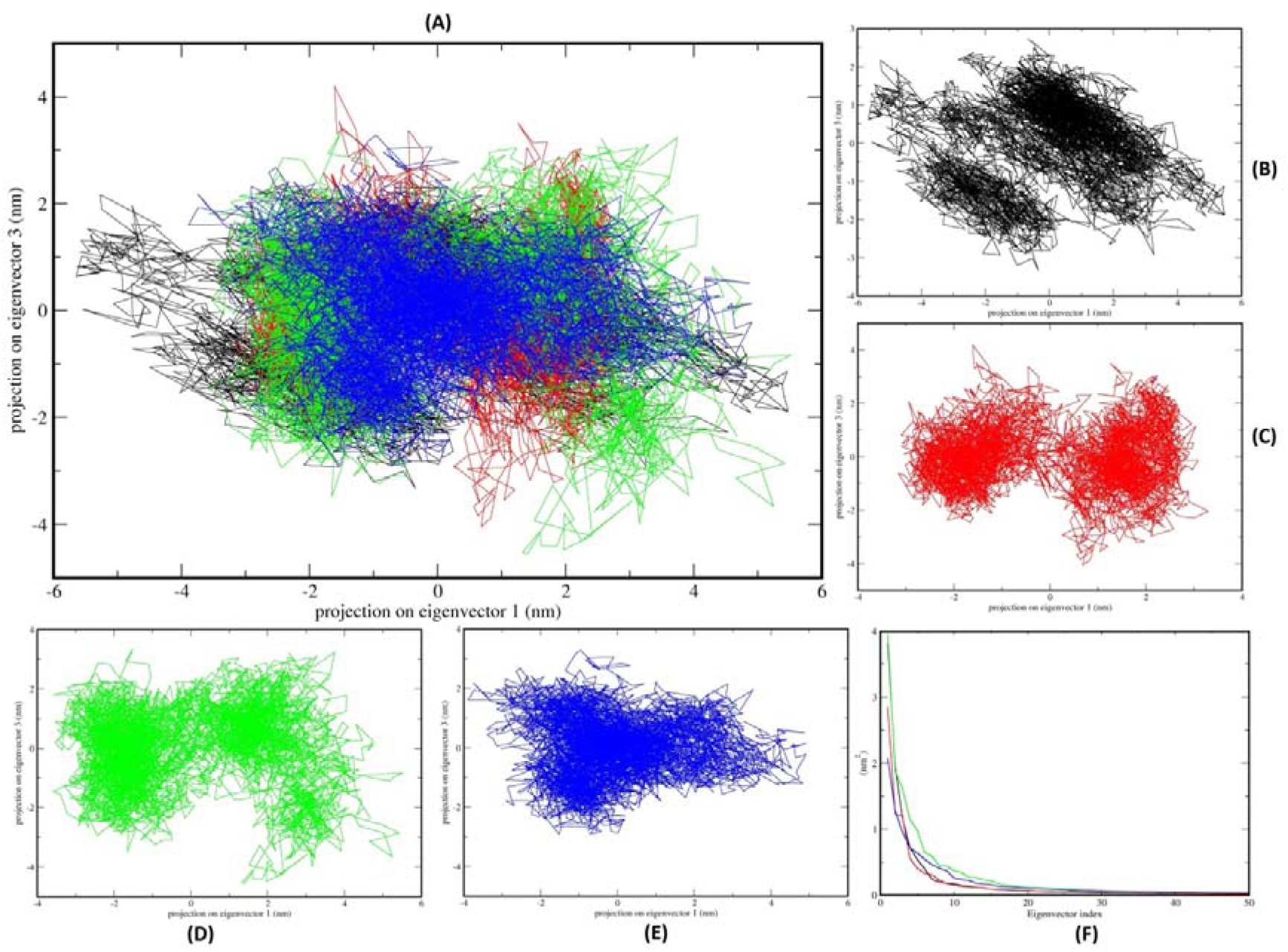
PCA 2D projection scatter plot of SARS-CoV and SARS-CoV-2 Mpro: (A) Overlay of 2D scatter plot projection the motion of the proteins in phase space for the two principle components, PC1 and PC3 derived from four MD simulation setup. Panel B, C, D, and E represent individual 2D plots of SARS-CoV_monomer, SARS-CoV-2_monomer, SARS-CoV_dimer, and SARS-CoV-2_dimer respectively. (F) Plot representing Eigenvalues calculated from the covariance matrix of Cα backbone fluctuations vs. the respective eigenvector indices for first 50 eigenvectors from 1000 eigenvectors. For all the panels colour representation is SARS-CoV_monomer (black), SARS-CoV-2_monomer (red), SARS-CoV_dimer (green) and SARS-CoV-2_dimer (blue).

To visualize the direction and extent of the principle motions, the first and last eigenvector was plotted in porcupine plot representation in which the arrows indicate the direction of eigenvector and magnitude of the corresponding value (Figure 14). From the plot, it was evident that SARS-CoV_monomer had cone projection throughout the three domains, while the SARS-CoV-2_monomer had less coherted motion in domain I and II. While the most stable and with least cone projection was the SARS-CoV-2_dimer indicating the most stable system formation. This result was is correlation was with PCA analysis

**Figure 14.**
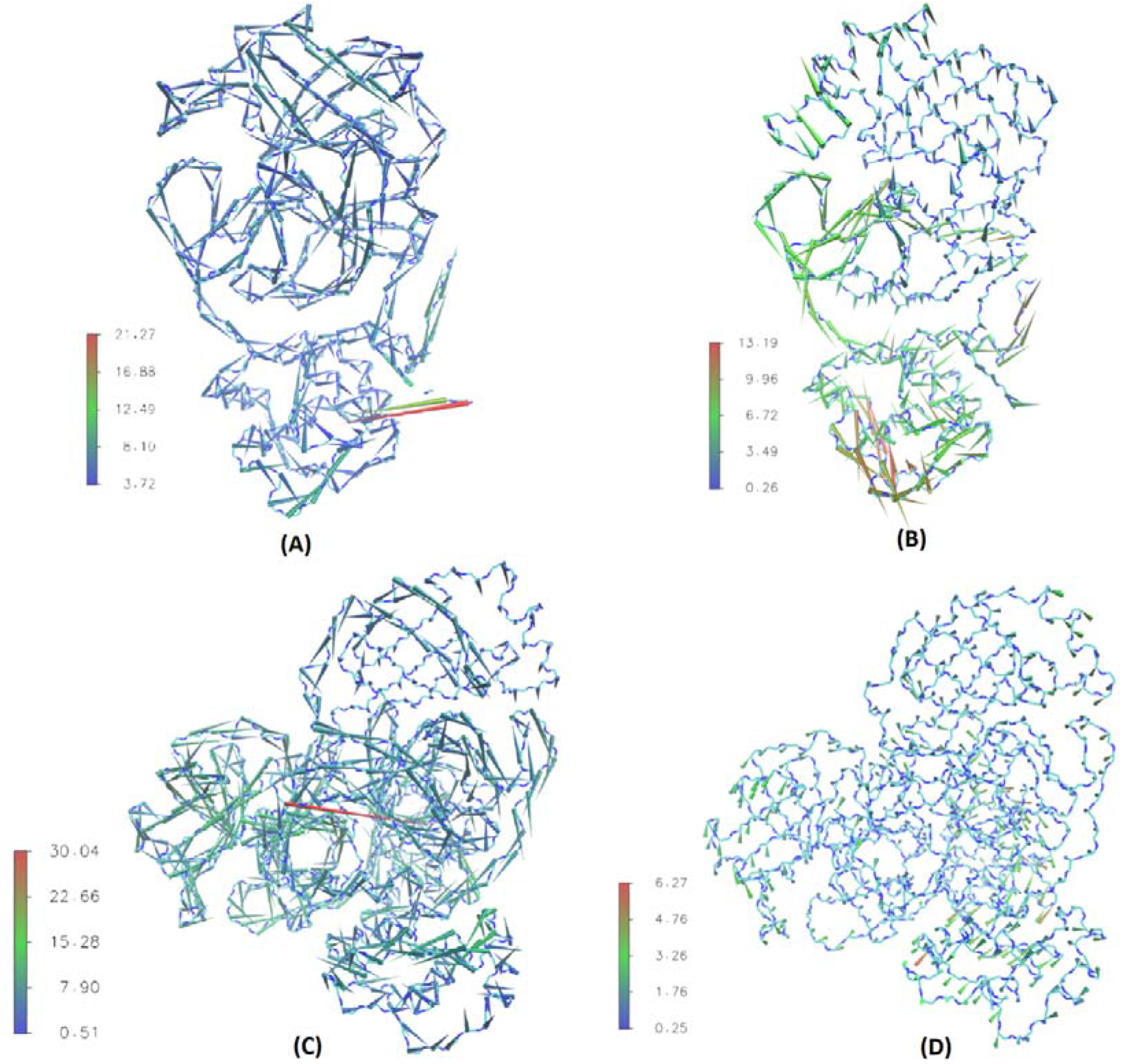
Porcupine plot for conformational variability computed from the crystal structure and average MD simulations ensemble. Porcupine plots of (A) SARS-CoV_monomer, (B) SARS-CoV-2_monomer, (C) SARS-CoV_dimer and (D) SARS-CoV-2_dimer. The length of the cone is proportional to the conformational variability, while the colour of the cone is represented by deviation in RMSD as indicated in the respective colour scale in each plot.

For generating representative structures for the conformational space traversed by the MDS, GROMOS [29] based clustering algorithm was applied on all four MDS setup. The method creates representative RMSD based clusters from the trajectory frames. It counts the number of the neighbouring structure using a 0.15 nm cut-off, and then form a cluster set with the largest numbers of neighbour structures followed by its elimination from the pool of clusters. The process is repeated for the remaining frames to identify other clusters with decreasing numbers of neighbour structures and each centroid of the cluster is used as a representative structure. These centroid structure members from each cluster are representative structures of distinct frames. The RMSD values in the clusters range from 0.0519-0.41 nm (average 0.158004) 0.0501-0.304 nm (average 0.146737), 0.0517-0.273 nm (average 0.139786) and 0.0532-0.247 nm (average 0.128409) for SARS-CoV_monomer, SARS-CoV-2_monomer, SARS-CoV_dimer and SARS-CoV-2_dimer respectively. A total of 19, 16, 12 and 5 clusters with 798, 1273, 637 and 158 transitions were observed for SARS-CoV_monomer, SARS-CoV-2_monomer, SARS-CoV_dimer, and SARS-CoV-2_dimer. The representative structures from each cluster from SARS-CoV_dimer, and SARS-CoV-2_dimer simulations are shown in supplementary Figure (S2, S3). Our cluster analysis indicated that the Mpro of SARS-CoV-2 in both the form is stable and less flexible than SARS-CoV Mpro, while the SARS-CoV-2_dimer had a stable and least number of conformations. The free energy surface analysis was performed for all the MD systems which represent the conformational variability in terms of ROG and RMSD taken together and represented by Gibbs free energy (Figure 15). The same trend as PCA and cluster analysis was also observed in FES. The both CoV_dimer Mpro had converged free energy and specifically, the SARS-CoV-2_dimer had the most converged free energy representing clustered RMSD and ROG values observed in its entire simulations.

**Figure 15.**
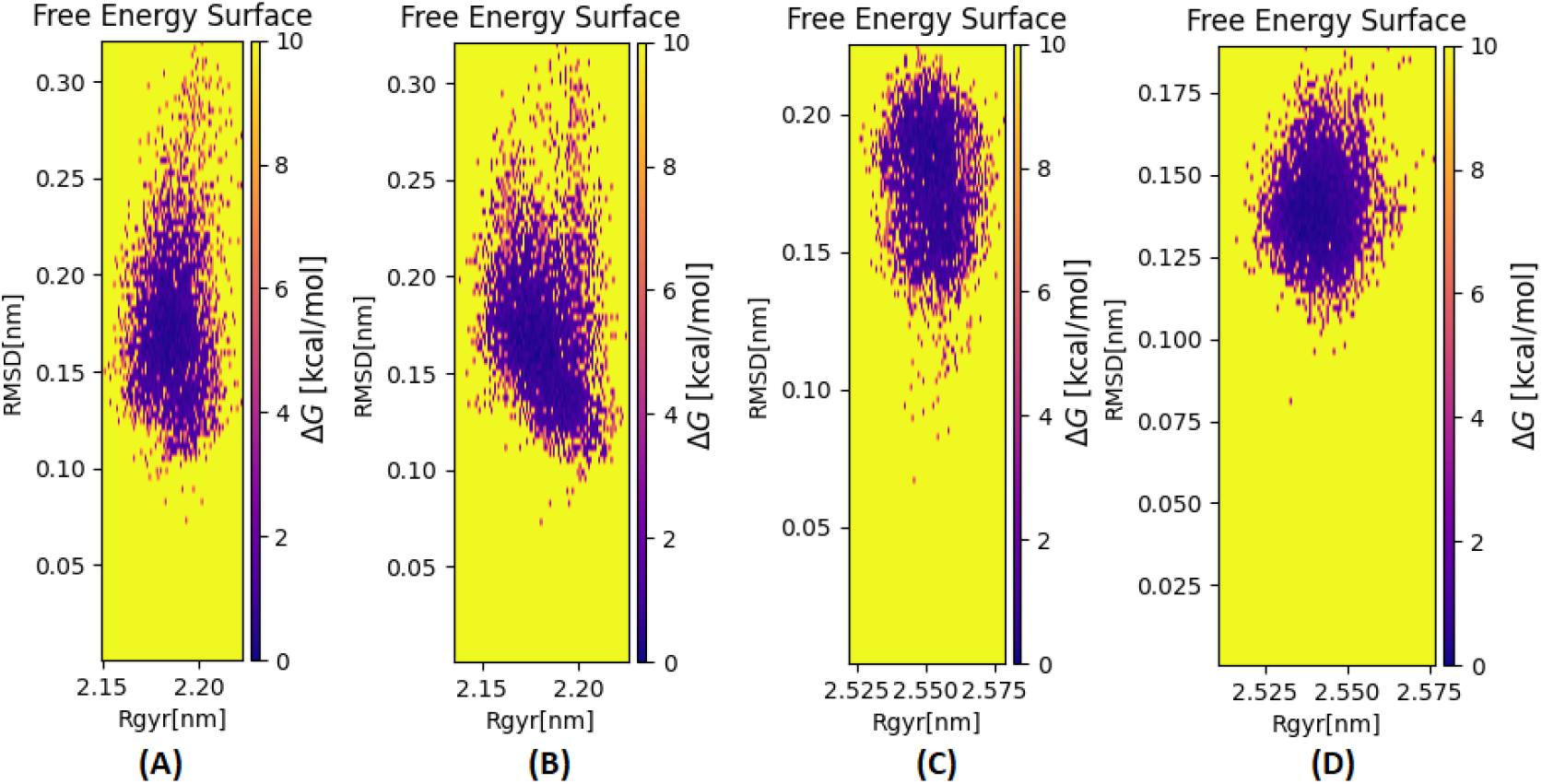
Free Energy Surface of Mpro computed over entire simulations (50ns): FES (in kcal/mol) for Mpro from (A) SARS-CoV_monomer, (B) SARS-CoV-2_monomer, (C) SARS-CoV_dimer and (D) SARS-CoV-2_dimer considering the conformational variability in terms of ROG and RMSD took together and represented by Gibbs free energy.

## Acknowledgement

Ritik and Meet acknowledges Institute of Advanced Research (IAR) for the resources. Dhaval Patel acknowledges the GSBTM, Govt. of Gujarat and Department of Biotechnology for DISC grant, Govt. of India for the resources and financial assistance. Prakash C Jha acknowledges Science and Engineering Research Board (SERB), Department of Science and Technology (DST) for a project grant, EMR/2016/003025. Mohd. Athar acknowledges generous support from the DST in the form of SRF INSPIRE Fellowship (IF150167)

## Supplementary Figures

**Figure S1.**
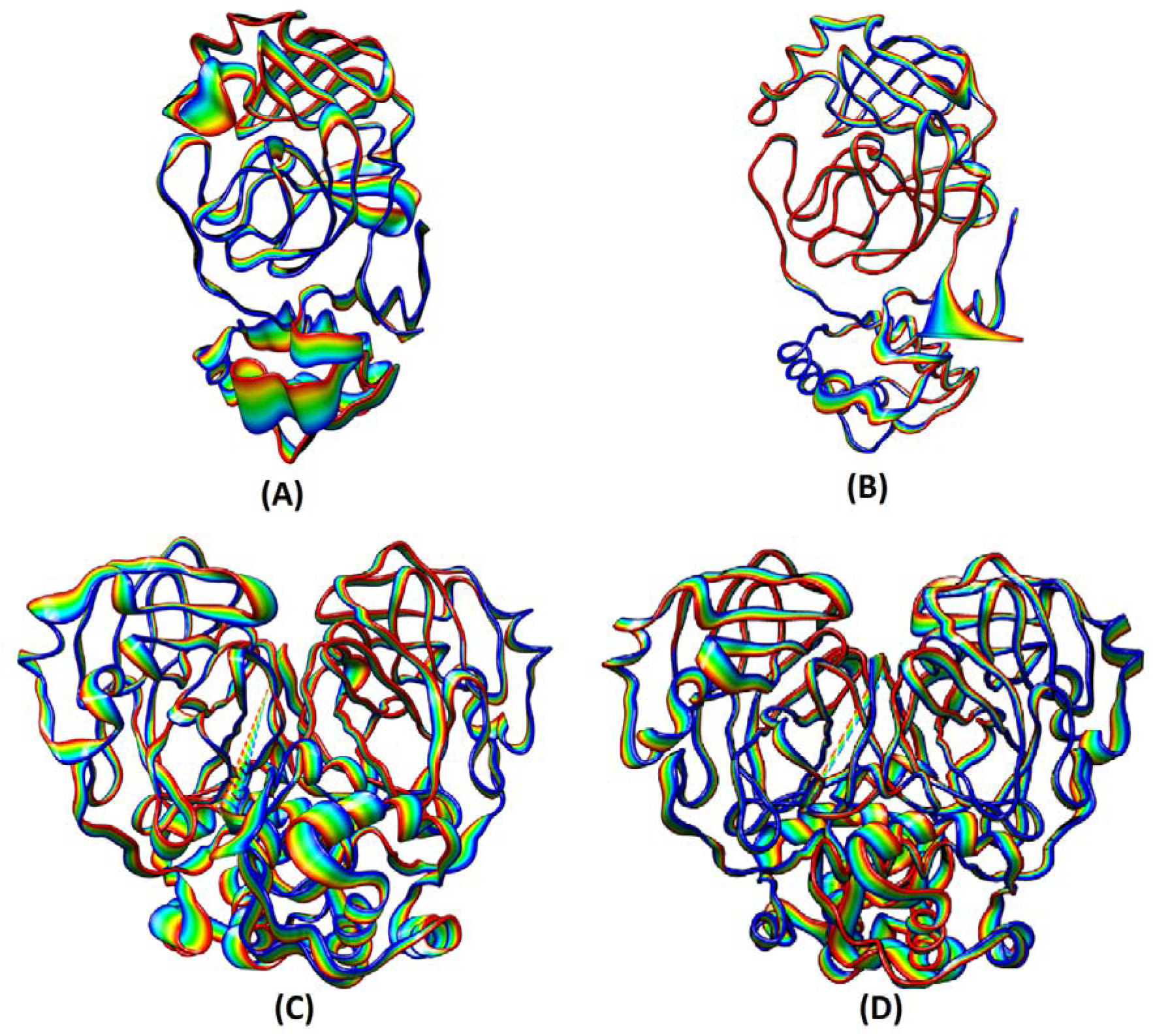
Comparison of 50 structures from each MD simulations projecting the extreme selected eigenvectors from PCA analysis for Mpro of (A) SARS-CoV_monomer, (B) SARS-CoV-2_monomer, SARS-CoV_dimer and (D) SARS-CoV-2_dimer. The width of main chain trace represents fluctuations throughout the timescale of MD simulations (50 ns) computed using PCA.

**Figure S2.**
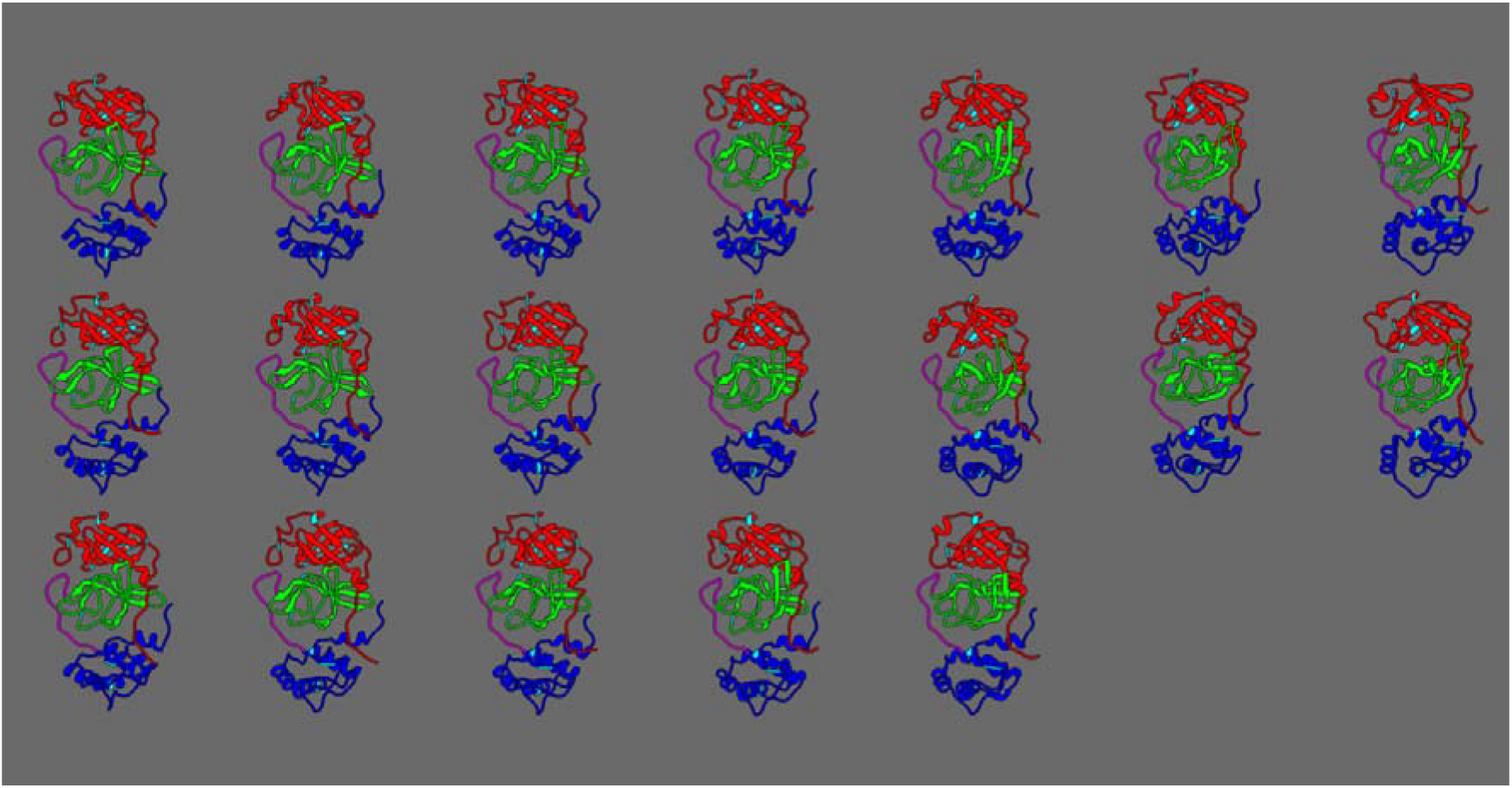
The representative structures from each cluster from SARS-CoV_dimer simulations. In all panel, the Domain-I is marked by red, Domain-II by green, Domain-III by blue and the loop region by magenta. The 12 variable residues are shown in cyan.

**Figure S3.**
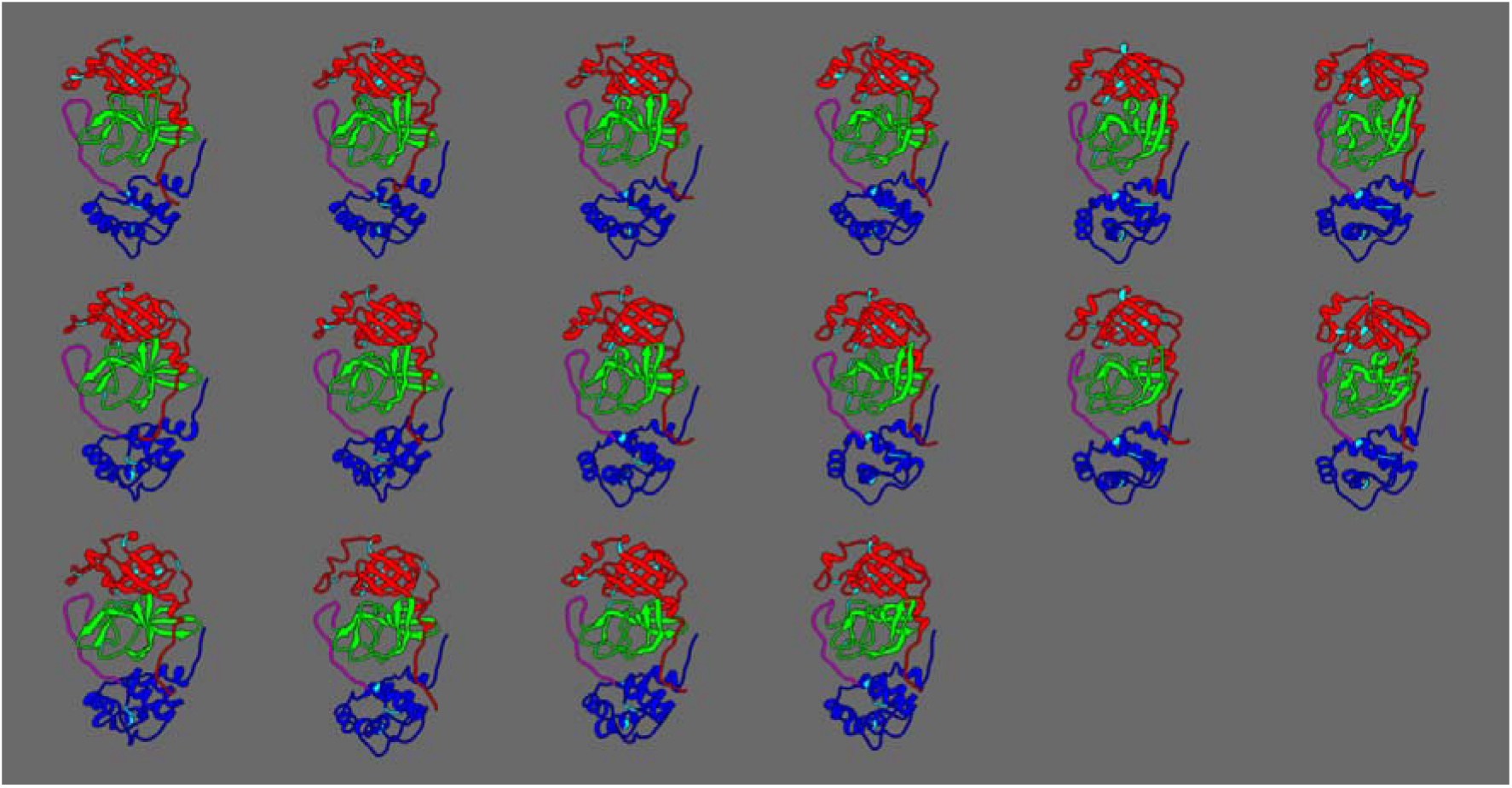
The representative structures from each cluster from SARS-CoV-2_dimer simulations. In all panel, the Domain-I is marked by red, Domain-II by green, Domain-III by blue and the loop region by magenta. The 12 variable residues are shown in cyan.

